# Machine learning guided design of strong transcriptional activation domains

**DOI:** 10.64898/2026.10.30.763198

**Authors:** Marissa A. Zintel, Claire J. LeBlanc, Sanjana R. Kotha, Melvin Soriano, Rishaan J. Bansal, Daniel Aguilera, Touss Majidi, Izaiah Ornelas, Liam D. Kirkpatrick, Aditya Udupa, Gean Hu, Pooja Agarwal, Angelica W. Y. Lam, James K. Nuñez, Patrick M. Shih, Rosangela Sozzani, Lucia C. Strader, Max V. Staller

## Abstract

Artificial intelligence has accelerated the design of structured proteins, but it remains difficult to design functional intrinsically disordered proteins because of weak structural constraints and poorly defined sequence-to-function relationships^1^. Activation domains (ADs), transcription factor regions that recruit coactivators, are an excellent test case for the *de novo* design of intrinsically disordered sequences. ADs are one of the oldest groups of functional intrinsically disordered sequences^2^ and designed ADs have applications in synthetic biology, biotechnology, and agriculture. We computationally generated 500,000 candidate sequences that resembled known ADs and predicted AD function with extant machine learning models, including three neural networks. From high-scoring sequences, we selected 18,750 for experimental validation in yeast. Surprisingly, >90% of the 16,770 well-measured sequences function as ADs. Sequences predicted to be active by multiple models were stronger, motivating development of a fast and accurate consensus AD predictor, MADRIGAL. A number of our designed ADs retain functionality across fungi, plants, and human cells. This work illustrates a generalizable paradigm for incorporating sequence-to-function models into the design loop to efficiently generate functional intrinsically disordered sequences.

## Introduction

The last few years have witnessed rapid progress in the ability to predict protein structure from protein sequence and design new structured proteins with specific functions^3,4^. This approach has been most successful when there is a strong coupling between structure and function^1^. Design of intrinsically disordered regions (IDRs) with cellular functionality has lagged behind because of weak sequence constraints and limited means to predict function from sequence^5–7^.

An ideal class of functional IDRs for *de novo* design are transcriptional ADs, the regions of transcription factors (TFs) that bind to coactivator complexes to activate gene expression. ADs were some of the first IDRs recognized to be functional^2^. Designed ADs have applications in synthetic biology^8^, animal models^9,10^, and biotechnology^11^. In agriculture, overexpression of TFs has created important GMO crops^12,13^, and designed ADs would expand this toolbox. Designed ADs would improve CRISPR activation methods to modulate gene expression, Perturb-seq, and epigenetic editing^14–17^.

Efforts to design ADs started nearly forty years ago^18^, but it remains challenging to design strong ADs. In screens of random peptides, 1-2% of sequences function as ADs^19–22^, but very few have high activity^23^. A larger fraction (up to 40%) of mixtures of aromatic and acidic residues can have high activity in yeast^20,24,25^, but not always in human cells^26^. Our past efforts to hand-design mutants to increase activity were successful in ∼⅓ of cases^26^. When we tested 18,000 sequences from the *S. cerevisiae* and *Arabidopsis thaliana* proteomes that had the strongest scores using the PADDLE neural-network predictor^27^, ⅔ had activity^28^. Although this success rate seems high, we expected all of the sequences to be active. Here, we test whether incorporating mechanistic insights and multiple predictors can improve our ability to design strong ADs.

Designing ADs is a stringent test of our understanding of the sequence features that control function. From high-throughput studies, it is clear that amino acid composition makes the largest contributions to activity and that amino acid arrangement, or sequence grammar, can also make large contributions to activity^24,25,27,29^. AD grammar is local and flexible: it includes clustering, intercalation, and balance between acidic and hydrophobic residues^24–26,30,31^. This balance is a critical part of our acidic exposure model, where acidic residues keep hydrophobic residues exposed to solvent and available to bind coactivators^32^. All known ADs that function across species and kingdoms are acidic ADs, the largest and most conserved known class^32,33^.

While it was historically difficult to predict ADs from protein sequence, recent high-throughput screens have trained neural network AD predictors^21,27,29,34^. While these predictors have high performance in cross-validation tests, they often generalize imperfectly to other datasets^28,31^. To compare the relative performance of existing AD predictors, we incorporated four predictors into our AD design and used two additional models in post hoc analysis.

We test the hypothesis that AD function can arise from the combination of composition and flexible local grammar that can be detected by neural network predictors. This hypothesis motivates a simple design strategy where we generate sequences with biased composition and local grammar using Markov chains, then select sequences with high scores from extant AD predictors for experimental testing. We found that >90% of tested sequences had AD activity in yeast. We found that sequences generated by longer Markov Chains were more likely to be predicted to be active, were more likely to be active in the experiment, and had stronger measured activity. Consensus between AD predictors identified the strongest designed ADs. This result motivated us to build a computationally efficient consensus model, MADRIGAL, for quickly predicting ADs. To prioritize candidates for synthetic plant biology and human cell CRISPR activation, we tested a set of medium and strong ADs in plants and human cell culture. We conclude that local grammar learned by Markov chains combined with neural networks efficiently designs strong ADs.

## Results

### Activation domain design

We designed strong ADs by generating sequences with increasing length-scales of local grammar and filtering them through four AD predictors. We titrated grammar by training Markov chains with state sizes between 1 and 5 residues to capture increasing local context (**Figure 1a**). The state size 1 Markov chain preserves only dipeptide frequencies, while the longest preserves enough context to incorporate short linear motifs of up to six residues (**Figure 1a**). Each Markov chain was trained on the same ∼6,200 strong plant ADs^34^, generating sequences with similar residue composition to known ADs. For each Markov chain, we generated 100,000 protein sequences, each with length 40AA, and prevented memorization by excluding sequences that had more than fifteen residues of overlap with any members of the training data (Methods, Table S1). The resulting sequences were very different from the yeast proteome, the training sequences, and each other, with the exception of clusters of sequences generated by the longest Markov chain (**Extended Data Figure 1a-b**). These clusters arise because the state size 5 Markov Chain has 290-fold more parameters than training data points (Methods). The designed sequences incorporate a range of sequence grammar based on Markov chain state sizes.

**Figure 1.**
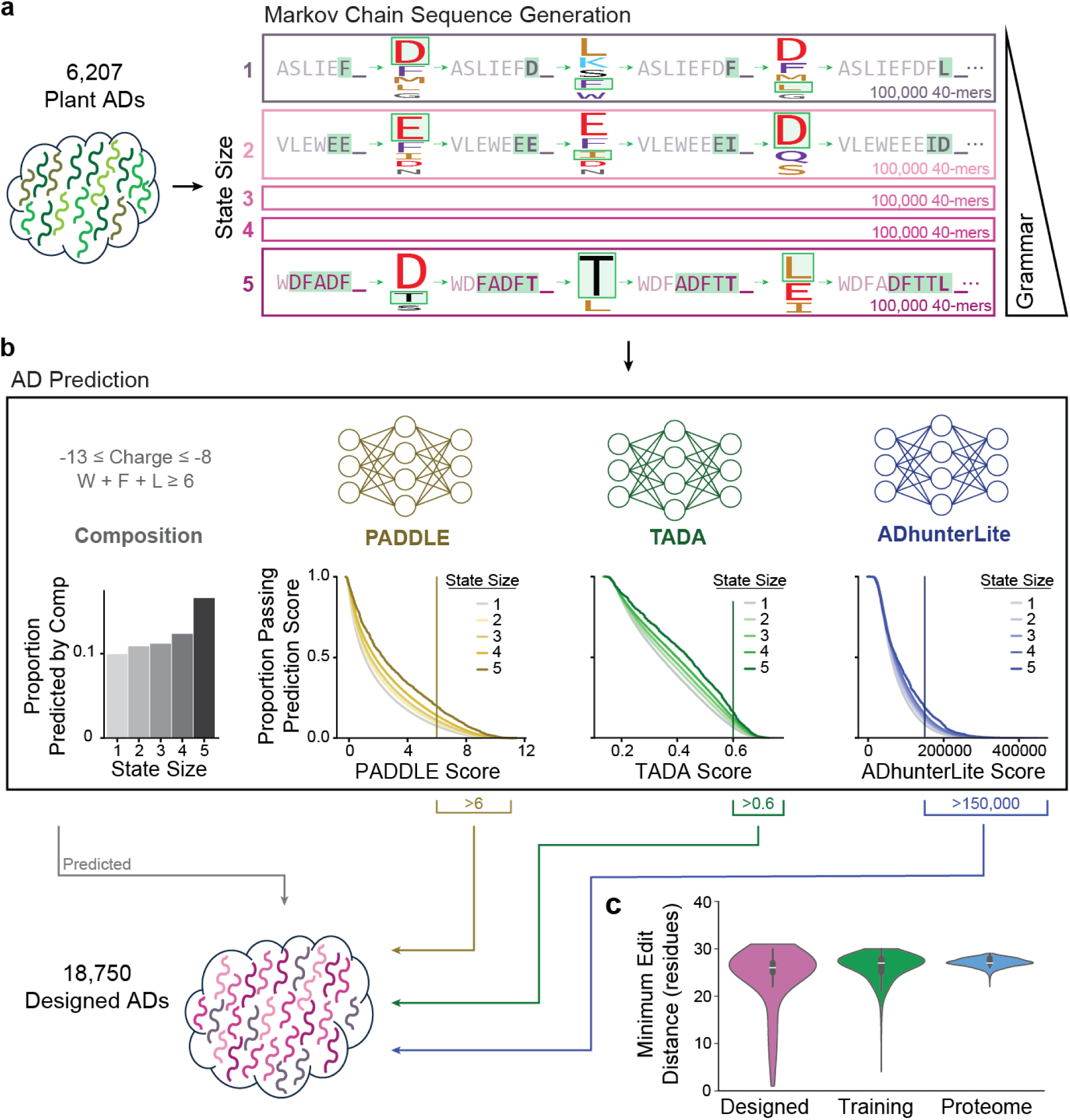
Pipeline for designing activation domains. a) 6,207 plant ADs were used to train Markov Models with state sizes 1-5. Increasing state size increases the local grammar incorporated into newly generated sequences. 100,000 sequences from each state size 1-5 were generated. b) All generated sequences were filtered through four predictors: Comp, PADDLE, TADA, and ADhunterLite. For Comp, the fraction of sequences predicted to be active is shown. For the neural networks, survival curves show the proportion of sequences predicted to be active as the threshold increases, with darker lines corresponding to higher state sizes. The vertical lines indicate the stringent thresholds used for selecting the 18,750 sequences. c) Minimum Levenshtein edit distance of the 18,750 experimental sequences to themselves (Designed), the training data, and the budding yeast proteome.

Sequences generated by longer Markov chains are predicted to be ADs at a higher rate because of both composition shifts and increased grammar. Mathematically, the longer Markov chains generate sequences with more grammar, but it is not a given that this will be the “right grammar” for AD function. To identify the strongest candidate ADs, we ran the Markov-chain-generated sequences through four AD predictors: Comp, PADDLE, TADA, and ADhunterLite (**Table 1**, **Figure 1b**). The Comp model is a “no grammar” control, while neural networks have learned some local grammar (Supplemental Note 1, **Extended Data Figure 2a-b**)^27,34–36^. Sequences generated from longer Markov chains had higher prediction scores, and were more frequently predicted active by the Comp model (**Figure 1b**). While composition contributes (**Extended Data Figure 1d-e**), the increase in prediction scores is primarily due to grammar, as shuffling and re-predicting the designed sequences lowered scores more for the higher state sizes (**Extended Data Figure 2c**). We conclude that longer Markov chain state sizes incorporate increasing amounts of sequence grammar that is recognized by the neural networks.

**Table 1:** Activation domain prediction models used in this study.

|  |  | **ADpred** | **PADDLE** | **Composition** | **TADA** | **ADhunterLite** | **ADhunter Ensemble** |
|---|---|---|---|---|---|---|---|
| **Input data** | Species of training data | Random peptides | *S. cerevisiae* TF tiles + mutants | human activation domains | *A. thaliana* TF tiles | Yeast Gcn4 homologs | *A. thaliana*, MycoCosm (fungal), and *S. cerevisiae* tiles |
|  | Measured in | *S. cerevisiae* | *S. cerevisiae* | Human | *S. cerevisiae* | *S. cerevisiae* | *S. cerevisiae* |
|  | Measurement | GFP | GFP (only kept cells within a narrow mCherry range) | Hand curated AD annotations | GFP:mCherry ratio | GFP:mCherry ratio | GFP:mCherry ratio |
|  | Size of dataset | 75,846 (10% test set) | 10,537 (~15% test set) | 888,581 39-mers, spaced 1AA | 70,937 (10% test set) | 18,947 (10% test set) | 25,660 (30% test set) |
|  | Positives in training set | 34,130 | 2,150 | 18,964 | 5,747 | 3,789 (top 20%) | 2,883 |
|  | Negatives in training set | 34,130 | 6,571 | 869,617 | 58,096 | 15,158 | 15,030 |
| **Model** | Input length | 30 AA | 53 AA | 39 AA | 40AA | 40AA | 53 AA |
|  | Features | Sequence | Sequence | AA counts | Sequence | Sequence | Sequence |
|  |  | Secondary structure | Secondary structure |  | 11 side-chain properties |  | ESM2 650M parameter model embeddings |
|  |  |  | Disorder |  | 9 disorder -related properties |  |  |
|  | Task | Classification | Regression | Classification | Classification | Regression | Regression |
|  | Architecture | convolutional | convolutional ensemble | simple composition | biLSTM | residual | residual ensemble |
|  | Parameters | 79,692 | 10 models, each with: 81,205 trainable and 81,879 total parameters | 7 | 430,736 | 96,769 | 14,716,436 for model plus 651,043,254 for ESM embedding |

### Nearly all designed activation domains function in yeast

For experimental testing, we selected 18,750 diverse sequences, with 3,750 from each Markov chain state size (**Figure 1c**, Methods; Table S1, Table S2). While model predictions were modestly correlated (**Extended Data Figure 3a**), we included some sequences where the predictors disagree to quantify relative model performance. We tested these candidate sequences with our Sort-seq high-throughput AD assay in *S. cerevisiae*^29^ (**Figure 2a-b**). We appended our candidates to the C-terminus of an inducible, mCherry-tagged synthetic TF, integrated them into yeast, and mated them to a GFP reporter strain. We sorted cells based on the ratio of GFP to mCherry, which detects the specific activity of each designed candidate AD (**Figure 2b)**. Our screen detected 16,770 designed sequences (89%), and the measurements had a large dynamic range and high reproducibility (Methods; **Extended Data Figure 4a**).

**Figure 2:**
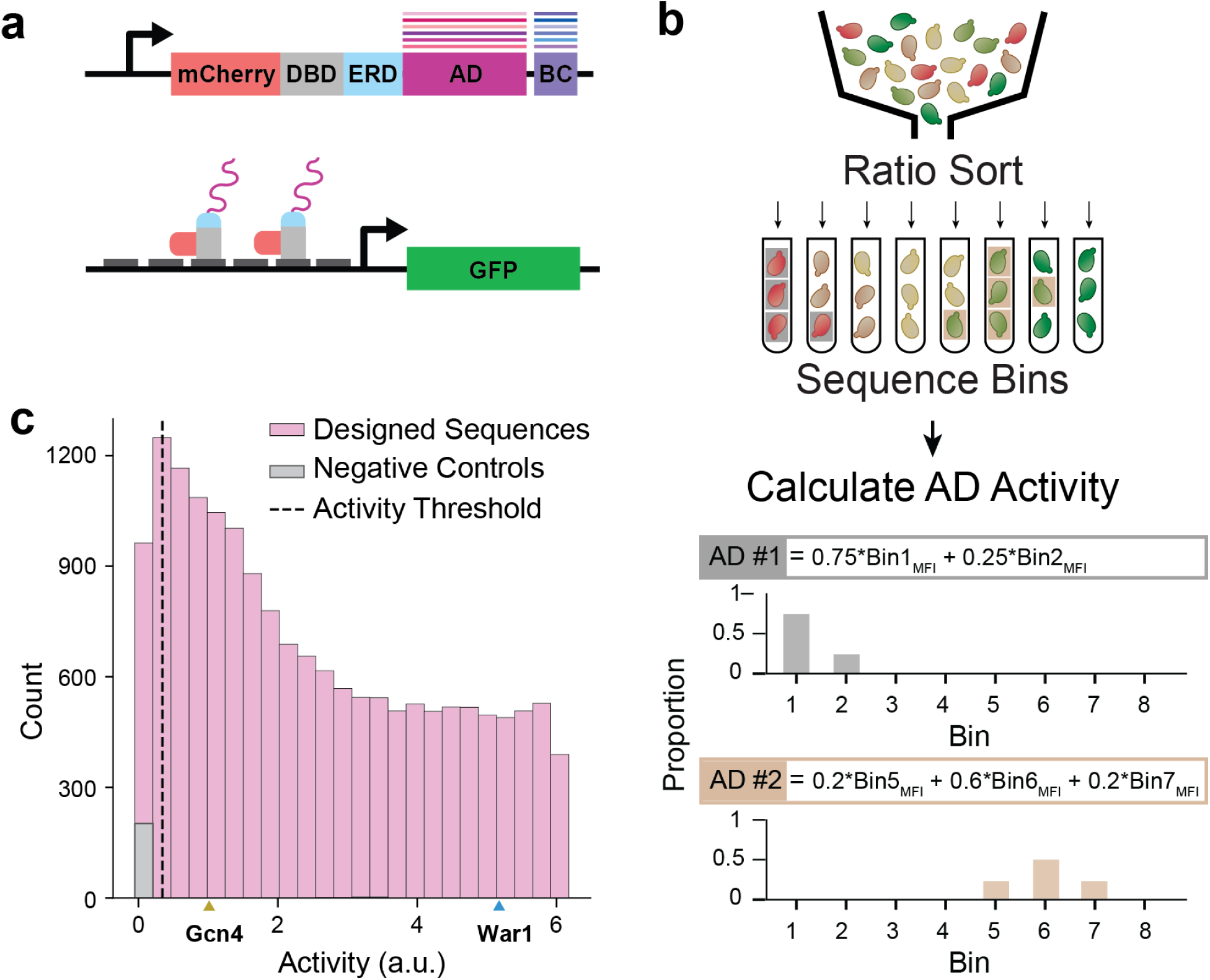
High-throughput experimental validation of designed activation domains in yeast. a) Synthetic TF construct and reporter construct, integrated on separate yeast chromosomes. DBD, DNA-binding domain; ERD, estrogen response domain; BC, barcode. b) Sort-seq assay workflow with GFP / mCherry ratio sorting and two examples of how activity is calculated as a weighted average. c) Activity distribution of all tested designed ADs, with the activities of two control ADs marked. The Gcn4 AD has an Activity of 1 and the Activity threshold is described in the Methods.

The combination of Markov chains and filtering with AD predictors efficiently generated strong ADs. 92.1% of the measured ADs had detectable activity **(Figure 2c**, Methods). Moreover, many of the designed ADs were very strong: 72.6% were more active than Gcn4 CAAD, a moderate AD; 11% were more active than the War1 C-terminal AD^29^, our strongest positive control from *S. cerevisiae*; and 2.9% were more active than CITED2, a very strong human AD. Our machine-learning-guided approach successfully designed many strong ADs.

The designed ADs have sequence features akin to extreme versions of natural acidic ADs. They have more aromatics, leucines, methionines, and more net negative charge than the training sequences (**Figure 3a-b**, **Extended Data Figure 5a**). The critical parameter is the balance between net charge and key hydrophobic residues: tryptophan, phenylalanine, tyrosine, leucine, and methionine (WFYLM, **Figure 3a**). While activity increases then decreases with respect to either net charge or WFYLM (**Figure 3b**), balanced increases in acidity and WFYLM content increase strength monotonically (**Figure 3a**). The requirement to balance net negative charge and WFYLM count supports our acidic exposure model^2,32^ (**Figure 3c**).

**Figure 3.**
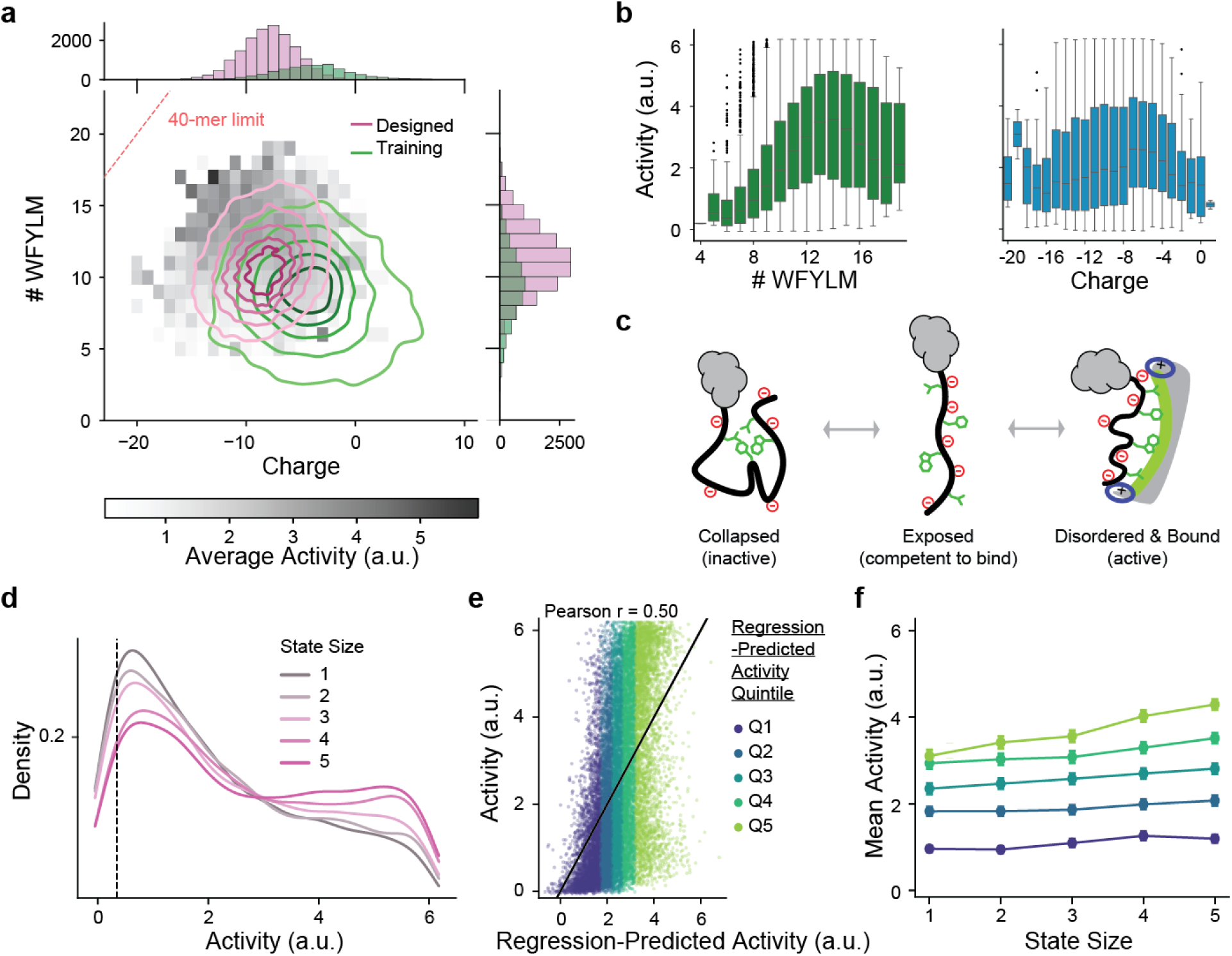
Strong activation domains balance acidic and hydrophobic residues. a) Distribution of hydrophobic residue composition (W/F/Y/L/M) and net charge for tested designed sequences vs. the 6,207 training sequences. Shaded boxes are the average activities of sequences within each composition box. b) Activity distributions by individual composition of hydrophobic and charged residues. c) Acidic exposure model^32^. d) Activity distributions by state size. e) Measured activity (y-axis) plotted against the predictions of composition-only linear regression (x-axis). The predicted activities are colored by quintiles. f) Grammar increases activity more for favorable compositions. Each line is the average activity of a quintile from composition-only linear regression predictions. Error bars are 95% confidence intervals.

Increasing sequence grammar boosts activity and synergizes with composition. Sequences generated from longer Markov chains were more likely to be active (**Extended Data Figure 5b**) and had higher activity (**Figure 3d**), even when controlling for composition (**Extended Data Figure 5c**). Compared to composition, the effect of grammar is modest. From state size 1 to state size 5, average activity increased by 25%, while from the bottom to the top quartile of WFYLM count, average activity increased by 167%. Yet, grammar and composition interact (**Extended Data Table 1**, Model 4 vs 3): sequences with more potent composition, as determined by linear regression on residue counts (**Figure 3e**), benefited more from longer grammar (**Figure 3f**). We conclude that once a sequence has the composition for high activity, grammar can amplify this potential.

### Consensus between predictors identifies the strongest designed ADs

For all neural network predictors, AD strength was correlated with prediction scores (**Figure 4b-f, Extended Data Figure 3b**). We assessed performance of the four predictors used in the design phase and added two more predictors, ADpred^21^ and ADhunterEnsemble^36^. Comp performs the worst (**Figure 4a, g, Extended Data Figure 3d**), while ADhunterLite and ADhunterEnsemble perform the best (**Figure 4e-g**, Table S1, Table S2). For a more detailed comparison of individual predictor performance, see Supplemental Note 1. Our analysis reemphasizes that the neural networks have high performance because they have learned sequence grammar.

**Figure 4.**
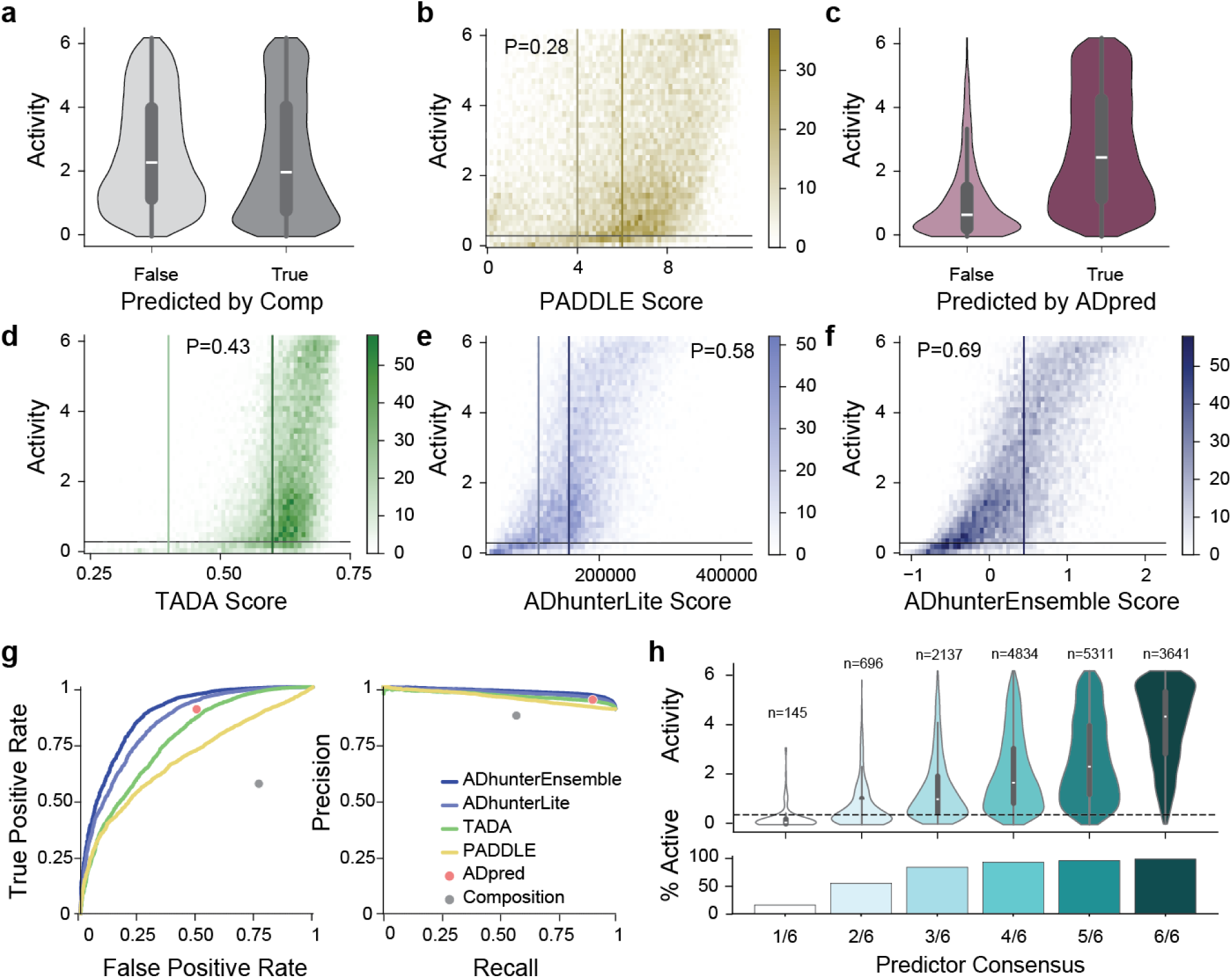
AD predictors’ performance on designed activation domains. Activities of designed sequences predicted by a) Comp, b) PADDLE, c) ADpred, d) TADA, e) ADhunterLite, f) ADhunterEnsemble. Pearson correlation coefficient is shown for models with quantitative output, and the standard and stringent thresholds are given by light and dark vertical lines, respectively. g) Receiver operator characteristic (ROC) and precision recall curve (PRC) plots showing classification performance of each predictor on the designed ADs dataset. True positives are defined as any sequences above the activity threshold, true negatives are defined as any sequences below the activity threshold. Scores for ADpred and Comp are shown as dots. h) Activity distributions and precision at increasing predictor consensus. Predictor consensus is the exact number of predictors (among a-f) that predict the set of sequences to be ADs.

Consensus between multiple AD predictors increased prediction precision (**Figure 4h**). Using standard published thresholds to binarize scores, each designed sequence was classified as an AD by 1-6 models (**Figure 4h**). Sequences predicted by only one or two models had low consensus, while sequences predicted by five or six models had high consensus. We found that increased consensus led to 1) a higher fraction of active sequences and 2) stronger ADs (**Figure 4h**). This trend was not driven by any single model or pair of models (**Extended Data Figure 3c**). Using the six-model consensus as a classifier had comparable performance to other neural network predictors (**Extended Data Figure 3e**). However, the designed sequences are very unbalanced, with more positives than typical datasets, so we set out to test model consensus as an AD predictor on natural TFs.

### A fast and accurate consensus activation domain predictor

Inspired by the increased precision of consensus between predictors, we built MADRIGAL, Multi-model Activation Domain Recurrent-neural-network Inferring Grouped Activity Labels. MADRIGAL is a surrogate model that is trained to predict the results of ensembling the six AD predictors. We collected over 200,000 unique, full length TF sequences from fungal, plant, and animal genomes, ran all six predictors to predict activity on tiling windows spanning all positions, thresholded the predictors, and trained MADRIGAL to predict the number of predictors that labeled each residue as active (**Figure 5a**, Methods). MADRIGAL is built with a long, short-term memory-based bidirectional recurrent neural network (LSTM-BRNN)^37^ trained and deployed using PARROT^38^. This enables MADRIGAL to take as an input a protein sequence of arbitrary length and output a per-residue prediction for model consensus (**Figure 5b**). With a Pearson correlation coefficient of 0.92 on sequences held out of training, MADRIGAL accurately captures the consensus knowledge of six heterogeneous AD predictors at a fraction of the computational cost (**Table 2**).

**Figure 5.**
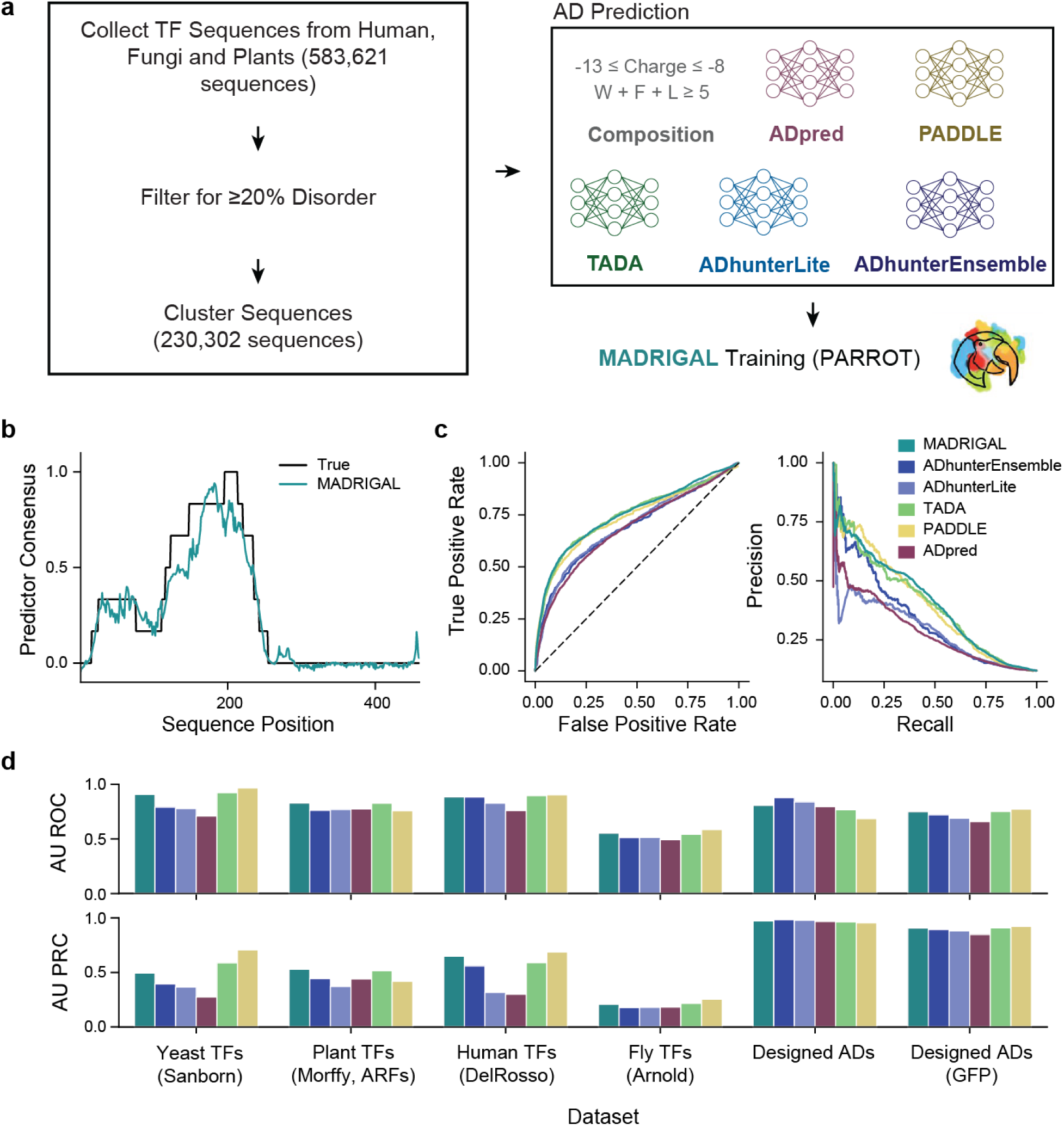
Design and benchmarking a Multi-model Activation Domain Recurrent-neural-network Inferring Grouped Activity Labels (MADRIGAL). a) To train MADRIGAL, disordered regions of eukaryotic TFs were collected and clustered by 75% sequence similarity. For 230,302 TFs, we ran all six predictors and then used PARROT^38^ to train MADRIGAL to mimic these predictions. b) MADRIGAL residue-level predictions in teal over an example sequence, with the actual six-predictor consensus in black. c) ROC and PRC plots for MADRIGAL and all neural network AD predictors on our entire test set of natural TFs (Methods). d) Area under the ROC and PR curve for MADRIGAL and individual predictors on each dataset separately. Sanborn and Morffy are measured in yeast, DelRosso in human cells, and Arnold in fly cell culture.

**Table 2:**
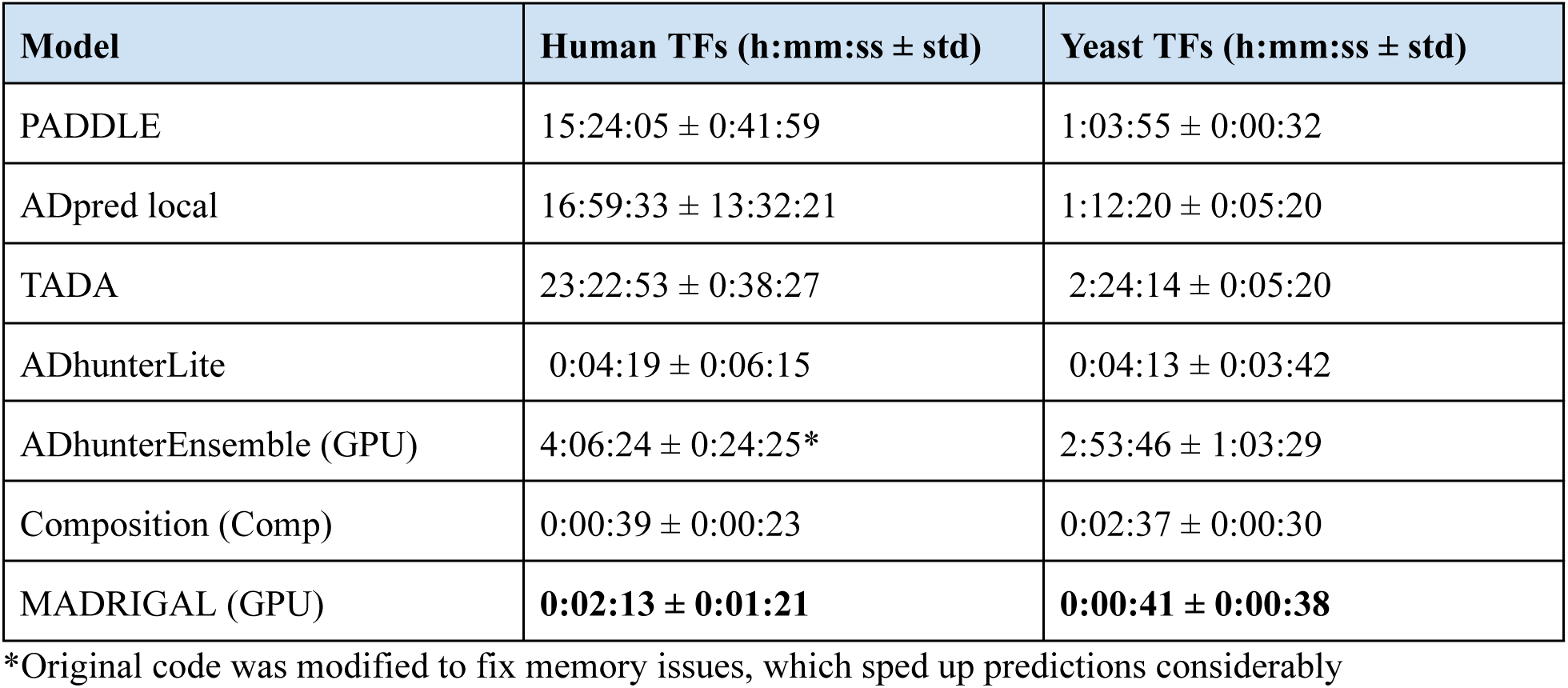
MADRIGAL model run times on human and yeast.

MADRIGAL rapidly predicts experimentally defined ADs with similar accuracy to existing AD predictors. We generated a test set of full-length proteins with known ADs, consisting of data from multiple independent studies that are absent from all predictors’ training sets (Methods). On our full test set, MADRIGAL is the best AD classifier by ROC and PRC AUCs (**Figure 5c**). Predictor performance varies by experimental dataset, and MADRIGAL is consistently one of the top three predictors within any experimental set, approaching the performance of the best predictor (**Figure 5d**, **Extended Data Figure 3f**). To evaluate how much each predictor contributed to the consensus predictor, we trained six leave-one-out models. All these models perform similarly to MADRIGAL, likely because of the high overlap between predictions (**Extended Data Figure 3a**). MADRIGAL has three advantages over individual predictors: 1) it is much more computationally efficient (**Table 2**), 2) it can run on arbitrary sequence lengths, making it simple to find ADs within TFs, and 3) it performs well across varied datasets.

### Testing designed ADs in other systems

To develop a toolkit of ADs for synthetic biology applications, we selected ten high-strength and four medium-strength ADs from our yeast screen (**Figure 6a-b**, Methods, **Extended Data Table 2**) to test in plant leaves and human cells.

**Figure 6.**
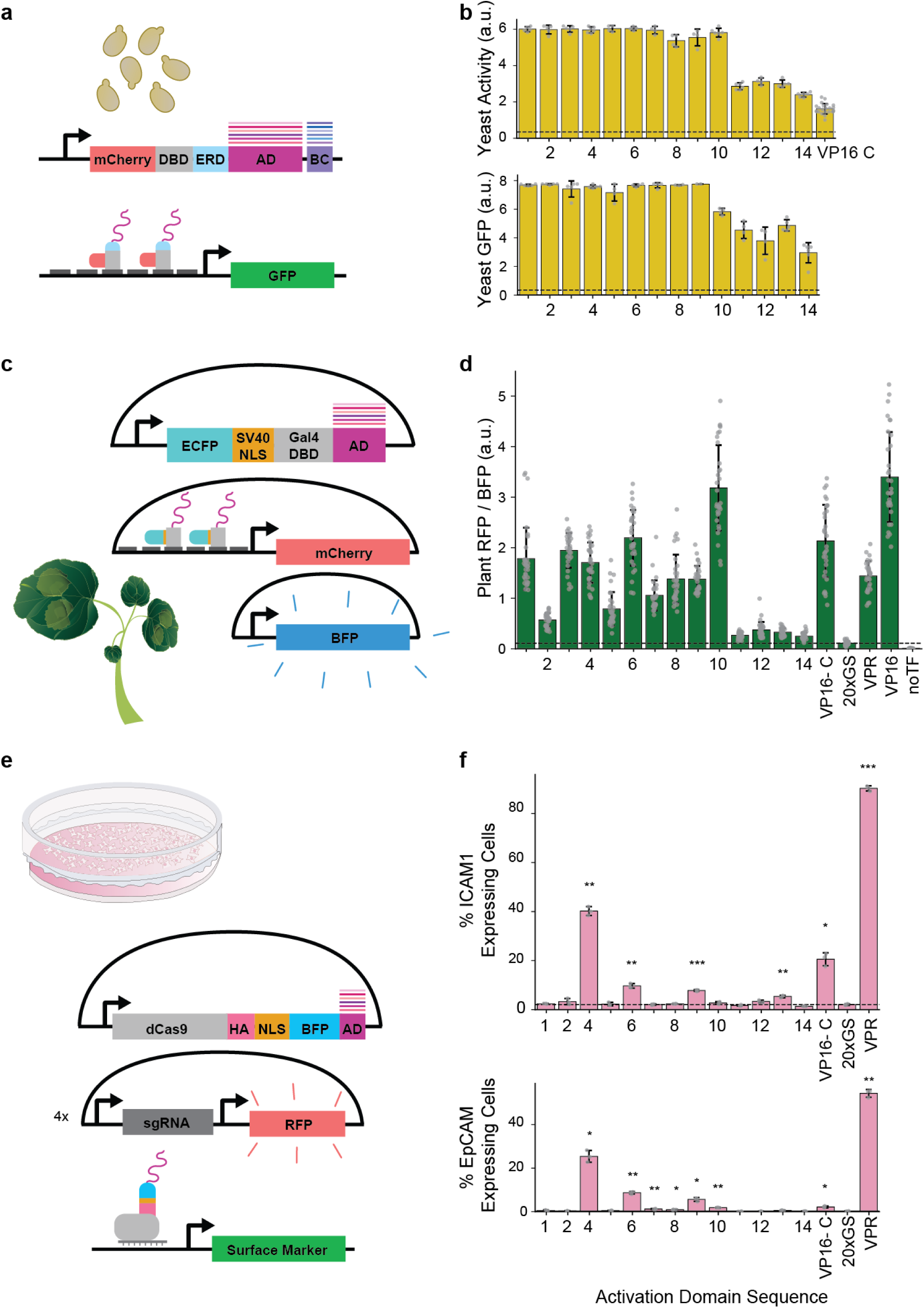
Designed ADs activate in plant and human cells. a) Yeast activator and reporter constructs. b) Average activity of each sequence in yeast sorted for AD activity (GFP / mCherry) or raw GFP. Error bars are standard deviations of 4-6 separately barcoded yeast (grey dots). c) Plant activator, reporter (mCherry), and transformation-normalizer (BFP) constructs. Each was delivered to plant cells via *Agrobacterium*-mediated transformation. d) Average activity of each sequence in plant leaves. Error bars are standard deviations of thirty-two measurements (four measurements per leaf, two leaves per plant, four plants). e) Transiently transfected CRISPRa activator, 4 sgRNAs, and reporter constructs target endogenous surface marker genes in HEK293T cells. f) Percent of cells expressing surface markers EpCAM or ICAM1 by antibody staining. Bars are the average of three transfections, error bars denote the standard deviation. Significance was calculated by Welch’s two-sided t-test with Benjamini-Hochberg correction within each plot, with replicates defined as 8 averaged plant leaves in (d) and 3 wells in (f) (*q<0.05, **q<0.01, ***q<0.001, all plant sequences were ***q<0.001).

To test AD activity in a plant context, we used *Agrobacterium* infiltrations (Methods) to transform *Nicotiana benthamiana* leaves (**Figure 6c**). We found that all fourteen designed ADs had significantly higher activity than a negative control GS-linker (**Figure 6d**). All ten of the high-activity sequences were more active than the four medium-activity sequences, a stronger preservation of rank order than we have previously seen^28,34,39^. Synth #10 was the strongest activator of plant leaf transcription and was indistinguishable from the VP-16 positive control despite being only forty amino acids instead of seventy-eight. These fourteen ADs encompass a range of activities that provide compact candidates for further development in plant synthetic biology.

To test activity in human cells, we used CRISPR activation (CRISPRa) targeted to endogenous cell surface marker genes in HEK293T cells (**Figure 6e**). Three of fourteen designed sequences clearly activated both EpCAM and ICAM1 (**Figure 6f**). Synth #4 was the strongest human cell activator, activating EpCAM and ICAM1 expression in 25% and 40% of cells, respectively – stronger than dCas9-VP16. Synth #4 activated weaker than the strong activator VPR (2.9-fold vs 8.7-fold for EpCAM and 2.4-fold vs 34-fold for ICAM1 over untransfected, (**Extended Data Figure 6a-b**)), although Synth #4 is a single 40 residue AD, while VPR is a tripartite 523 residue chimera. Synths #6 and #9 also activated both EpCAM and ICAM1, albeit at lower levels. Our machine-learning-guided design yielded at least three ADs (#4, 6, 9) that function across three kingdoms of life: fungi, plants, and animal cells.

## Discussion

Leveraging recent advances in our understanding of AD function, we combine local grammar and composition encoded by Markov chains with neural network predictors of ADs to efficiently design very strong ADs. Our designed ADs compositionally resemble extreme versions of the natural acidic ADs explained by our acidic exposure model: they contain clusters of hydrophobic residues balanced by an acidic context.

Our systematic titration of grammar reinforces the flexible role of local grammar. Given a favorable composition, high activity can arise without grammar^20,21,24^. However, for a given composition, differential patterning can boost or depress activity (**Figure 3f**). We find that longer grammar (at least up to six residues) boosts activity, that the clustering and intercalation of hydrophobic and acidic residues is important, and that the grammar is complementary, but secondary, to composition. Our results are consistent with a recently proposed hierarchy of grammar rules^24,25^. All the neural network AD predictors have learned aspects of this grammar.

Consensus between predictors identified the strongest ADs, implying that the different predictors are learning orthogonal information. This result motivated us to create an ensemble-surrogate predictor, MADRIGAL, to combine learned information from six independent training datasets. MADRIGAL is computationally efficient, makes length-agnostic, residue-level predictions, and has strong performance on varied datasets. Because the best AD predictor varies by dataset, MADRIGAL is ideal for quickly screening sequences to identify candidate ADs or querying TF evolution in large families. We created a website (https://www.madrigal-predictor.net/) and a Python package for running MADRIGAL (https://pypi.org/project/madrigal-predictor/).

Our machine-learning-guided design method has yielded novel ADs that activate across fungal, plant, and animal kingdoms of life. All fourteen designed ADs tested in *N. benthamiana* leaves successfully activated a plant leaf reporter, encompassing a broad range of activities. Synth #10 activated as well as the VP-16 AD, one of the strongest activators in the system^28,40^. In HEK293T cell CRISPRa fusions, three ADs activated two endogenous human genes, a more difficult task than activating a reporter with multiple TF binding sites^15,16,41,42^. Consistent with prior studies, there is substantial plant- and human-specific variation in AD strength^28,34,43^. These designed ADs are compact, orthogonal sequences that span a broad range of activities in plants and include activators of endogenous human genes, providing foundational parts for a synthetic gene activation toolkit across eukaryotic kingdoms.

AD prediction and design are currently dominated by acidic ADs. Acidic ADs are generally the strongest ADs in high-throughput datasets and, as a result, all current AD neural network predictors reserve their highest prediction scores for acidic ADs. We analyzed predicted ADs from thousands of diverse TFs and found that more stringent thresholds shifted the composition of predicted ADs towards more negative charge and higher WFYLM counts for every neural network predictor (**Extended Data Figure 7**). It is currently unclear whether the lower AD prediction scores for non-acidic ADs arise because the predictors have learned that non-acidic ADs have weaker activity or if the predictors are blind to other AD classes.

Using ADs as a test case for designing functional IDRs, we demonstrate that a simple pipeline incorporating biased composition, local grammar, and neural network predictors can efficiently generate strong ADs. For acidic ADs in yeast, we have achieved predictable design at scale.

## Methods

### Design pipeline for generating activation domains

To generate candidate activation domains, we trained a Markov chain sequence generator (markovify, https://github.com/jsvine/markovify) with the 6,207 active sequences (activity > 1) from a tiling screen of *Arabidopsis thaliana* TF sequences in yeast (the plant activation domain identification (PADI) dataset)^34^. Markovify generates new protein sequences (sentences) residue-by-residue based on the amino acid (word) sequence patterns in the training data. We used Markov chains with state sizes 1-5 to generate 40-residue sequences based on context windows of 1-5 residues. For each state size, we generated 100,000 sequences with at most 15 residues allowed to overlap exactly with any sequence from the training data. After performing the experiments, we determined that the state size 5 model was under-constrained, because there were more potential parameters (64M) in the model than there were 6mers in the training data.

All 500,000 generated sequences were then predicted by four activation domain predictors: Comp, PADDLE, TADA, and ADhunterLite. We used Comp ^31^ as described in the literature. We used PADDLE without secondary structure ^27^ with a threshold of 6, which is more stringent than the standard threshold of 4. Because PADDLE takes 53-mer inputs, we appended the constant 13 amino acids upstream of the designed activation domain to calculate prediction scores. For design, we used an early version of TADA^34^, and a threshold of 0.6, which is more stringent than the recommended threshold of 0.4. ADhunterlite (originally named ADhunter_v1 in^36^ but retrained in^30^) is a regression model, and we used 150,000 as a stringent threshold for design. For downstream analysis we used recommended thresholds, and PADDLE with secondary structure. See Supp Note 1 for comparisons of model performance.

Within the 100,000 sequences generated from each state size, we intended to select sequences from every combination of agreement among the four predictors (i.e. 4/4 models agreed, all four versions of 3/4 agreeing, eight versions of 2/4 agreeing, and 4 versions of only one model predicting a sequence to be active). We discarded sequences predicted by none of the models and sorted the remaining sequences into one of 15 categories based on which combination of the four predictors predicted those sequences as ADs. However, categories were defined non-exclusively, so many categories that were intended to be predicted by only 1, 2, or 3 models were in fact predicted by more. We drew 250 sequences from each category for each state, totalling 18,750 sequences, with 3,750 from each state size.

We also computed mean pairwise Levenshtein distances for every sequence to all other generated sequences within each category. These distances were >30 for all sequences, indicating that sequences were overall diverse. When choosing each set of 250 sequences, we selected sequences with the lowest mean pairwise distances in their category (combination of predictors, non-exclusive), collecting the most typical sequences per category. The minimum pairwise distances remain high among the final sequences overall (most greater than 20). The exception was state size 5 because it is data-limited, so it contains clusters of similar sequences. There are no duplicate designed sequences.

Our final set of generated sequences is 18,750 diverse synthetic 40-mers that are very different from both the training sequences and each other.

### Yeast library generation

We ordered the 18,750 designed ADs plus 204 inactive, negative control sequences as a Twist oligo pool (Twist Biosciences). We reverse translated designed amino acid sequences to DNA using optimal yeast codons. Polynucleotide runs and restriction enzyme sites were removed by synonymous mutations.

The plasmid library was generated as previously described^30^ with the addition of random barcodes to distinguish between distinct yeast integration clones. The pMVS219 vector (also called pMVS142; Addgene 99049) was digested overnight with NheI-HF, PacI, and AscI, heat inactivated at 80C, run on a 0.8% agarose gel, and gel purified (NEB^®^). The oligo library was 18,954 sequences with 436.2 ng yield. This pool was resuspended in 100 µL water. The oligo pool was PCR-amplified (Q5) for 6 cycles with LC3.P1 and YL_randBCs_R3_3’ primers to add random 14 bp barcodes. Eight PCR reactions were pooled into two PCR clean up columns (NEB^®^). The amplicon was cloned into the C-terminus of the synthetic TF with Gibson assembly using NEB^®^ HiFi Mix (4 reactions). The assemblies were cleaned up on a PCR column (NEB^®^) before electroporation into NEB^®^ 10-beta electrocompetent cells.

The plasmid library was digested with SalI-HF and EcoRI-HF restriction enzymes (NEB^®^) to generate a fragment containing the synthetic TF and the KANMX cassette. This fragment was transformed into the DHY211 yeast strain (MY566) yeast cells along with two amplicons of flanking sequences with homology to the URA3 locus. The upstream fragment bridges URA3 with the ACT1 promoter (pMVS295 with YP18 and CP19.P6 for 626 bp), while the downstream fragment bridges the KANMX terminator and URA3 (pMVS296 with YP7 and YP19 for 665 bp). Transformants were grown overnight on YPD and then replica plated onto YPD+G418 (200 µg/ml). The integrated yeast library was mated to MY435 (FY5, MATɑ, YBR032w::P3 GFP ClonNat-R (pMVS102)) and diploids were selected in liquid YPD with 200 µg/ml G418 and 100 µg/ml ClonNat.

We also constructed a Controls Library to normalize across previous Sort-seq experiments and impose a consistent threshold for activity (Table S3). The Controls Library contained 497 sequences from LeBlanc et al., 2026^30^: 200 negative control sequences, 100 strong activation domains, and 197 activation domains spanning intermediate strengths. The Controls Library allows for very accurate calibration and harmonization of Sort-seq experiments.

### Yeast Sort-seq experiment

We sorted yeast cells as previously described^30^. Briefly, designed ADs and Controls yeast libraries were grown overnight, separately, from 100 µL of a glycerol stock. On the day of sorting, the controls yeast library was spiked into the designed ADs library as 5% of the cells based on OD_600_. Yeast were then diluted 1:5 in SC-Glucose + 1 μM β-estradiol to induce nuclear localization of the synthetic TF, and incubated at 30°C for 4 hours. Yeast were concentrated by centrifuging at 3,000xg for five minutes, filtered through a 35 µm membrane and stored on ice until sorting.

Yeast cells were sorted using a BD Aria Fusion five times: a GFP sort (raw reporter activity per cell), an mCherry sort (TF abundance per cell), and three GFP/mCherry sorts (AD “Activity”: reporter activity normalized by TF abundance) on two separate days. Yeast were sorted into 4 or 8 bins, with each bin containing approximately one million cells in all bins. Bin boundaries were drawn to separate the fluorescence distributions into approximately equally populated bins (**Extended Data Table 3**). In our first sort (Sort 1), we sorted on GFP with 8 bins, mCherry with 4 bins, and GFP/mCherry twice, with 8 and 4 bins. After collecting additional yeast colonies, we sorted the designed library by GFP/mCherry ratio with 8 bins again, this time with the control library spiked-in (Sort 2). In GFP/mCherry ratio sorting, it is easy to saturate the FACS detectors ^30^, so we carefully calibrated the fluorescence of the bins to maximize dynamic range and reduce saturation. This designed AD library had the highest activity we have ever measured, spreading out positive controls that had previously clustered in the top bin.

After sorting, fresh SC-Glucose media was added to each bin up to ∼6mL, and cells from each bin were grown overnight at 30°C. The next day, cells were pelleted and gDNA extracted using the Zymo YeaStar Genomic DNA Kit (D2002, Protocol I with chloroform). Barcoded AD sequences were amplified from 100 ng of gDNA per bin as previously described^30^ starting with CP21.P14 and CP17.P12.

### Sequencing and activity calculation

Sequencing libraries were generated from the first PCR with two more steps: first using 4 ng of template and a pool of SL5.F1-4 and SL5.R1-4 phasing primers for 6 cycles with Q5^®^ (NEB^®^, cat M0492) and second using MVS_00XX_I2 and MVS_00XX_I1 for 6 cycles, where “00XX” indicates unique index pairs for NovaSeq compatibility. Libraries were sequenced by 150PE reads, on 2 shared lanes of the NovaSeq X 1.5B platform for Sort 1 and 1 lane of the NovaSeq X 10B platform for Sort 2. Reads were paired using PEAR^44^, then used as inputs to our published Sort-seq activity calculation functions (https://github.com/staller-lab/labtools). Reads were normalized first by the total reads in each bin, second by the number of cells in each bin, and finally across the eight bins to calculate the relative abundance of each barcode across the eighth populations. These relative abundances across each bin were multiplied by the bin median fluorescence intensity as weighted average, or dot product. We observed high technical reproducibility between sorts (**Extended Data Figure 4a**): barcode measurements agreed strongly (**Extended Data Figure 4b**) indicating high biological reproducibility. Distinct barcodes were averaged together after discarding outlier measurements (>2.6 standard deviation, 50,553 barcodes included and 361 barcodes excluded for Sort 2). Sort 2 had more barcodes (50,553 vs 43,332 in Sort 1, Ratio) and was measured with the Controls Library spiked-in, so we used Sort 2 in all Activity analyses.

### Using controls library to harmonize with LeBlanc et al., 2026

After activity calculations, the negative controls and Controls Library sequences were used to harmonize the data to LeBlanc et al., 2026. First, activity scores for all controls were correlated between this work’s Sort 2 and LeBlanc et al., 2026. Scores were linearly correlated at medium and low activities, but LeBlanc et al., 2026 scores saturated earlier than scores measured in this work. Thus, we normalized the two datasets based on the least active sequences and a moderate AD, Gcn4. A Gaussian curve was fit to the negative control sequences, and the activity at its peak (20268 in LeBlanc et al., 2026; 5390 in this work) was set to 0, while Gcn4’s activity (106674; 46954, respectively) was set to 1. Because the GFP sort did not include the controls library, we used the mean activity of 20 designed sequences that were very similar to the activity of Gcn4 (as measured in Sort 2), as the Gcn4 “1” Activity for all reporter sorts. LeBlanc et al., 2026 set their activity threshold based on 0.01 FWER across all sequences in their library, with a score corresponding to 0.28 in Gcn4-normalized units. mCherry scores are raw FACS values (a.u.).

### Protein sequence analyses

Sequence features Net_Charge, Kappa, Omega, etc. were calculated for each designed sequence using localCIDER (https://github.com/Pappulab/localCIDER)^45^. Sequences were also predicted post-hoc by ADpred^21^ and ADhunterEnsemble^36^. ADpred computes per-residue scores and predicts an activation domain if at least 10 consecutive residues score at least 0.8. We use ADpred primarily as a binary predictor, but we also evaluate the mean of per-residue scores to evaluate ADpred as a continuous predictor. ADhunterEnsemble, like PADDLE, takes 53-mer input sequences, so we similarly appended the constant 13 amino acids upstream of the designed AD to calculate prediction scores. ADhunterEnsemble is intended as a regression model, not for classification, but in order to binarize its scores to measure predictor consensus, we used the classification-proxy threshold of 0.44 (Waldburger et al., 2026; dataset mean + 1 standard deviation).

We used statsmodels to fit four linear regression models of activity by ordinary least squares. Model 1 (composition only) used counts of all 20 amino acids as numeric variables; Model 2 (state size only) used state size as a numeric variable; Model 3 (composition + state size) used all 20 amino acid counts and state size as variables; and Model 4 (composition + state size interaction) used all 20 amino acid counts plus all counts with a state size interaction term. State size was centered at 1. Models 1, 3, and 4 did not use intercepts because all sequences have 40 residues, meaning residue counts will always sum to 40 and provide an intercept term. Similarly, Model 4 did not include state size alone because the per-residue state size interaction coefficients sum to 40 x state size. For each model, we used five-fold cross validation. Because state size 5 contains many similar sequences, we clustered sequences based on 50% identity (MMseqs2) and used these clusters to partition sequences into the five folds using GroupKFold (sklearn). Standard errors for model coefficients were likewise clustered, and p-values were corrected via the Benjamini-Hochberg procedure within each model. R^2^ was computed from out-of-fold predictions as 1 - SSE/SST. Using the full dataset, we evaluated each model by F-tests against an intercept-only model, and also evaluated Model 3 vs 1 and Model 4 vs 3, to test the significance of increasingly complex models. As a further clustering control, we also recreated all 4 models on two additional splits of the data to confirm the state size effects: a dataset of the representative sequences from clustering and a dataset restricted to state sizes 1-4 (**Extended Data Table 1**).

To control for the largest composition differences across state sizes with an orthogonal approach to linear regression, we first calculated the most variable residues across state sizes. For all measured designed sequences (unclustered), we calculated the mean amino acid count within each state size, and used the range of the five state size means to quantify the total variation for each residue across state sizes. We then binned the most variable residues into quantiles, ensuring that each composition stratum had at least 10 sequences present in each of at least 3 state sizes. The binned residues and their bin counts were chosen by progressively lowering a residual threshold and checking that both the binned and un-binned residues varied by less than the residual threshold, while ensuring that at least 80% of sequences were retained. This resulted in binning S composition into 7 quantiles, and L, D, G, A, E into 2 quantiles, retaining 89.5% of sequences in 117 distinct composition strata. Within each stratum, each residue’s mean count varies by at most 0.2 across state sizes. We then quantified mean activity per state size by subtracting each strata’s overall mean from the mean of each state size (within a strata), and averaged these for all strata. This results in deviations from the mean Activity for each state size, so the mean Activity was added to each state size deviation for plotting comparisons.

To determine the impact of SLiMs on activity, we first searched the experimentally measured sequences for three previously reported motifs: WxxLF, LxxLL, and ΦxxΦΦ, where Φ is any hydrophobic residue W, F, Y, L, M, I, or V. WxxLF was present in only 0.4% (67/16770) sequences, so was not included in subsequent analyses. To control for hydrophobic residue content, we stratified on leucine count or total Φ count for LxxLL and ΦxxΦΦ analyses, respectively, keeping only strata with at least 10 sequences with and without the motif. We then compared motif-containing to composition-matched sequences by Welch’s t-test with Benjamini-Hochberg correction across strata.

### Assessing the Grammar Detection of Neural Network AD Predictors

To confirm that the neural networks have learned AD grammar signals, we ran the five neural network predictors on two shuffle datasets. Staller et al., 2018^29^ measured the activity of Gcn4 and >900 compositionally-identical shuffled versions^29^. Sanborn et al., 2021^27^ measured the activity of 8 yeast ADs and 8 shuffles of each. We calculated Pearson correlations for each shuffle set. To determine if the grammar incorporated by Markov chains was recognized by AD predictors, we shuffled all generated sequences once using NumPy’s default random number generator, seed=20261008. We then re-predicted each shuffled sequence.

### Prediction metrics

Receiver operating characteristic (ROC) and precision-recall (PR) curves were generated with sklearn. Areas under the curves were calculated by trapezoidal area. Sequences above 0.28 were considered as active (positives); sequences below 0.28 were considered inactive (negatives) for both GFP/mCherry and GFP scores.

We evaluated the predictors, including MADRIGAL, on variable length TFs with identified activation domains from other experiments. We denoted true activation domains as defined in the original experiments. Specifically, for yeast sequences, we used ADs in “Figure 1—source data 2” from Sanborn et al.^27^. We only used ADs found in TFs that were held out of PADDLE’s training data (as defined in “Figure 3—source data 1”). For plant sequences, we used “Supplemental Table 3” from Morffy et al.^34^. For human sequences, we used “Supplementary Table 2” from Delrosso et al.^46^. For fly sequences, we used “Table EV2” from Arnold et al.^23^. Any residue that was part of an activation domain by the original paper was considered a “true” activation domain residue. All residues that were predicted to be part of an activation domain were considered “predicted” activation domain residues. For the predictors that required fixed length inputs, we tiled each full length sequence with a step size of 1 residue and then ran the predictors on all tiles. Any residue within a tile that was predicted to be active was considered to be a “predicted” activation domain residue. From our set of true and predicted residues, we calculated true positives, false positives, true negatives, and false negatives and used these values to construct ROCs and PRCs.

Using these prediction definitions, we assessed the compositional preferences of the predictors at standard and stringent thresholds. Across the 230,302 TF sequences for which we predicted ADs (see below), the Net Charge and WFYLM count were calculated for predicted vs background sequences. For each window-based predictor, three random 40-mers or 53-mers were sampled for every predicted contiguous AD. Net Charge and WFYLM count were averaged across all random samples. 53-mer samples were normalized to 40-mer counts by multiplying by 40/53. For ADpred, the “standard” criteria defined ADs as at least 10 consecutive residues passing the 0.8 score threshold, and each sample was the 40-mer centered on a randomly chosen ≥0.8 residue. The “stringent” criteria was defined as requiring the entire sampled 40-mer to be within a stretch of ≥0.8 scores. Background composition was calculated by sampling three random 40-mers per protein sequence, and averaging the composition counts.

### Training MADRIGAL

We collected 310,656 fungal TFs, 172,854 Plant TFs, and 137,436 metazoan TFs. We filtered sequences on length (53-1442 AA), filtered out sequences with less than 20% predicted disorder (Metapredict2^47^), and clustered sequences with at least 75% sequence similarity, yielding 230,302 sequences. This clustering prevents overlap between sequences, which will help avoid overfitting. We ran all 6 predictors on windows that tiled across the entire length of all 230,302 sequences. We found that 1.96% of residues are predicted by four or more predictors, and 3.91% are predicted by three or more predictors. We binarized the predictions of each predictor and trained a PARROT model on the number of positive predictors at each position.

The PARROT architecture consists of two bi-directional long short-term memory (biLSTM) layers and dense output layer^38^. To validate the performance, we built two test sets. One had 18,388 independently measured AD sequences from multiple publications^20,23,27,28,34,46^. In Figure 5c, performance was assessed on an independent validation set of 16 Yeast TFs with 12,981 residues, 49 Fly TFs with 33,772 residues, 355 Plant TFs with 101,141 residues, and 25 Human TFs with 13,895 residues. We added the 16,770 designed activation domains measured here as a separate validation set, because they were not included in the training data. MADRIGAL can be run on the website https://www.madrigal-predictor.net/ or the package can be downloaded and run locally from https://pypi.org/project/madrigal-predictor/.

### Benchmarking the run times of different predictors

For every model, we ran the prediction 3 times (n=3). For most models, we ran on savio3 using a single node with 32 Intel Xeon Skylake 6230 CPUs. We used the original prediction code (without any modifications to speed them up). The only exceptions are ADhunterEnsemble and MADRIGAL (because they are designed to work with GPUs). We used savio3 along with a single NVIDIA GeForce RTX 2080 Ti GPU.

### Selection of activation domains for plant and human experiments

To test the universality of our designed activation domains, we selected 10 high activity and 4 medium activity designed ADs from the yeast screens to test in both plant and human cell assays. We selected the 10 high activity sequences (1-10) by picking the highest scoring sequences in the GFP/mCherry sort with at least four replicates (separately barcoded instances of the designed AD) that agreed well. We also selected 4 (11-14) medium activity sequences using the same procedure, to assess rank order of activities across systems and potentially define a toolkit of varying strength ADs. We also verified that these sequences were measured in both the GFP and mCherry sorts, and ensured that mCherry (abundance) scores of the selected ADs were not low, to mitigate low expression issues in plant and human cell systems.

After selecting the 14 sequences, the sequences were ordered as oligo pools along with a negative control flexible linker sequence (20xGS) and the C-terminal 40 amino acids of the VP-16 activation domain (VP16-C), which was a control sequence with low-moderate activity in the yeast GFP/mCherry sort. Plant and human cell assays used the 20xGS as a negative control AD sequence and each assay’s individual highest activity activation domain as a positive control.

### Plant experiments

*In planta* activity was measured via a *Nicotiana benthamiana* transient transformation assay utilizing fluorescent protein reporters. Designed AD and control sequences were cloned into a synthetic TF construct within the T-DNA of a binary vector plasmid (pAS89). These synthetic TFs were expressed from a constitutive promoter (pCL1)^28^ and contained an N-terminal CFP tag, Gal4 DNA binding domain, and the AD of interest at the C-terminus. Plasmids were transformed into electrocompetent *Agrobacterium* strain GV3101. A reporter T-DNA binary vector (pAS61) containing 6x Gal4 UAS and a 35S minimal promoter driving mCherry expression, and a normalizer T-DNA binary vector (pAS40) with the constitutive pMAS promoter driving BFP expression were also transformed into GV3101. Transformed cells were selected on LB agar plates containing kanamycin (50 µg/mL), gentamicin (30 µg/mL), and rifampicin (100 µg/mL). Single colonies were cultured overnight in liquid LB with the same antibiotics and refreshed the day of infiltration by 1:5 dilution. After 2.5 hours of growth, cultures of the TF, reporter, and normalizer *Agrobacterium* strains were pelleted, resuspended in infiltration buffer (10 mM MgCl2, 10 mM MES pH 5.6, 200 µM acetosyringone), and mixed in a 1:1:1 ratio to a final OD_600_ of 0.133 per strain, followed by a 1-hour induction at room temperature. Each TF / reporter / normalizer *Agrobacterium* mixture was then infiltrated into 2 leaves (4^th^ and 5^th^ from the top) of 4 4-week-old *Nicotiana benthamiana* plants. 4 days post-infiltration, 4 6-mm leaf discs were extracted from each infiltrated leaf and placed abaxial side up atop 350 µL of water in black, clear-bottom, 96-well microtiter plates (Corning). mCherry and BFP fluorescence from each leaf disc was measured using a BioTek Synergy H1 microplate reader (Agilent) with excitation/emission wavelengths of 587nm/615nm and 400nm/454nm respectively. To determine the activity of each AD, mCherry fluorescence from the 6x Gal4 UAS reporter was normalized to BFP fluorescence from the constitutively expressed normalizer.

### HEK293T CRISPRa Assay

Activities of dCas9-fusions were measured by transient transfection and recruitment to two endogenous gene targets, EpCAM and ICAM1. HEK293T cells were maintained under standard culture conditions (DMEM supplemented with 10% (v/v) FBS (VWR) and 100 U/mL streptomycin, 100 mg/mL penicillin (Gibco)) and seeded into 96-well plates at 15,000 cells per well one day prior to transfection to enable 3 biological replicate measurements for each dCas9-fusion and target gene combination. Cells were co-transfected with 150 ng of dCas9-effector plasmid and 75 ng of 4-pooled sgRNA plasmids targeting either the endogenous EpCAM or ICAM1 loci. The dCas9-effector plasmid consists of catalytically inactive Cas9 (dCas9), an HA tag, nuclear localization sequence, blue fluorescent protein (BFP), and the indicated effector domain. sgRNA expression plasmids contain an mCherry reporter to enable identification of sgRNA-positive cells. Two days post-transfection, cells were harvested and stained for surface expression of commercially available FITC labeled antibodies for EpCAM or ICAM1 (BioLegend) following the manufacturer’s protocol.

Cells were analyzed by flow cytometry using BD FACSymphony A1 Cell Analyzer (BD Biosciences) flow cytometer. BFP, mCherry, and GFP fluorescence were measured to identify cells with the dCas9-effector fusion, sgRNA, and target protein expression, respectively. Cells were first gated for single, live cells, followed by selection of BFP-positive and mCherry-positive populations corresponding to successful delivery of both dCas9-effector and sgRNA components. Effector activity was quantified by measuring target gene activation within the BFP-positive/mCherry-positive population using both GFP median fluorescence intensity (MFI) and the percentage of GFP-positive cells. Gating strategy for all three untransfected replicates is shown in **Extended Data Figure 6c-d**. Measurements from untransfected control cells were calculated from the total live, single-cell population and used as baseline expression controls.

### Use of LLMs

LLMs Claude and Gemini were used for some code generation with close supervision.

## Data availability

All the raw sequencing data are available in GEO under accession number GSE352640. All generated sequences and predictions are in Table S1. The candidate activation domains selected for the experiment, their predicted activities, and their measured activities are in Table S2. Predictions of the 6 AD models for the 230,302 sequences are available in the github.

## Code availability

Github for analysis and MADRIGAL development: https://github.com/staller-lab/Madrigal-development, 10.5281/zenodo.23813869

The MADRIGAL package is available at https://pypi.org/project/madrigal-predictor/

Github for sequence analysis: https://github.com/staller-lab/designed-ads.git, 10.5281/zenodo.23816789

## Limitations of this study

Our designed ADs are generated to resemble known acidic ADs, which constrains the explored sequence space. Like the vast majority of natural ADs, we do not know what coactivators our designed ADs are recruiting. In contrast, the co-submitted manuscript by Chen et al. designed structured proteins to bind to individual coactivators^48^. Based on work by others^27^, we suspect our designed activation domains are binding to Mediator. The existing AD predictors remain biased towards predicting acidic ADs, and, consequently, our method can only design acidic ADs. Moreover, all the existing neural networks are yeast predictors. Even TADA, which excels at finding plant ADs, is trained on plant sequences measured in yeast. These predictors can find acidic ADs but struggle with other classes. The field now has a mature understanding of the sequence features and grammar that control the function of acidic/aromatic ADs. This group appears to be the oldest and largest class in eukaryotes, with members working well across distantly related species. Future studies with new reporter systems in other species are needed to identify, predict, and eventually design other classes of ADs, including glutamine-rich, proline-rich, and serine-rich ADs.

## Author Contributions

Conceptualization, Funding Acquisition: LCS, RS, MVS

Data Curation: MAZ, CJL, MS, SRK

Formal Analysis: MAZ, CJL

Software: MAZ, SRK, CJL, MS, RJB, GJH, PA, AWYL

Investigation: MAZ, MS, DA, TM, IO, LK, AU

Supervision: JKN, PMS, LCS, RS, MVS

Visualization: MAZ, MVS

Writing–original draft: MAZ, MVS

Writing–review & editing: MAZ, CJL, JKN, PMS, LCS, RS, MVS

## Acknowledgements

We thank Nick Ingolia, Alex Holehouse, Ryan Emenecker, Gloria Brar, Alex Chavez, Zeba Wunderlich and members of the Staller Lab for comments on the manuscript. The DHY211 yeast strain was a gift from Angela Chu and Joe Horecka.

## Supplementary Information is available for this paper

Correspondence and requests for materials should be addressed to MVS.

