## Extended Data Figures for "Machine learning guided design of strong transcriptional activation domains"

**Table of contents**

**Extended Data Figure 1. Sequence properties of the candidate designed ADs. 1**

**Extended Data Figure 2. The neural network AD predictors recognize AD grammar. 2**

**Extended Data Figure 3. AD predictor analyses. 3**

**Extended Data Figure 4: Assessment of Sort-seq measurement quality. 4**

**Extended Data Figure 5. Relationship between composition and AD activity. 5**

**Extended Data Figure 6. Fluorescence flow cytometry on human cells. 6**

**Extended Data Figure 7. Sequences predicted to be active under more stringent prediction thresholds are more negatively charged and have more hydrophobic residues. 7**

**References 7**

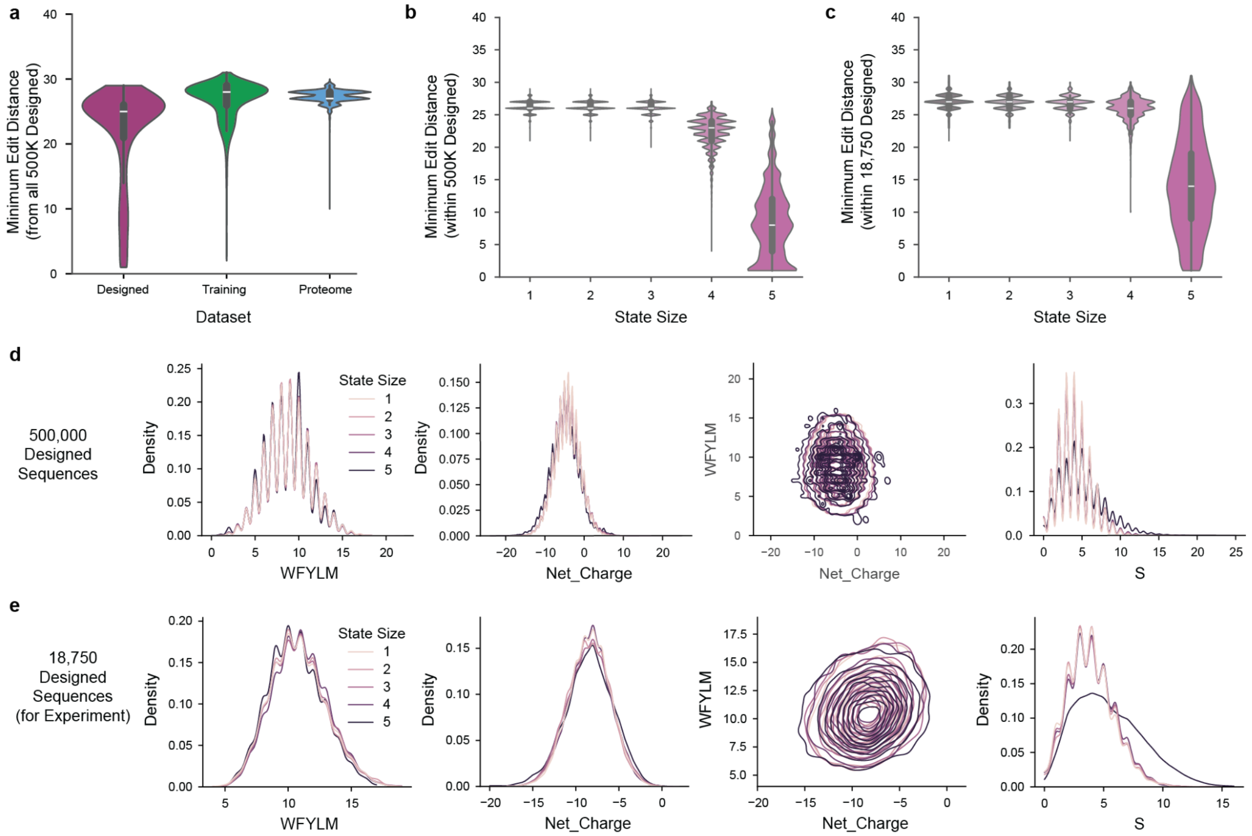


**Extended Data Figure 1. Sequence properties of the candidate designed ADs.**
a) The distribution of minimum edit distances of every generated sequence (500,000) to every other generated sequence, the training dataset, and the budding yeast proteome. b) The distribution of minimum edit distances within all generated sequences, as in (a), but split by state size. c) As in (b), but for only the 18,750 sequences selected for experimental testing. For all the generated sequences (d) and the experimental sequences (e) the distribution of net charge and the number of W,F,Y,L, and M residues is consistent across the five state sizes. The combination of these two parameters, which are important for AD strength, is consistent across state sizes. The single amino acid with the largest difference between state sizes is serine (S): the state size 5 Markov chain generated more sequences with more serines.

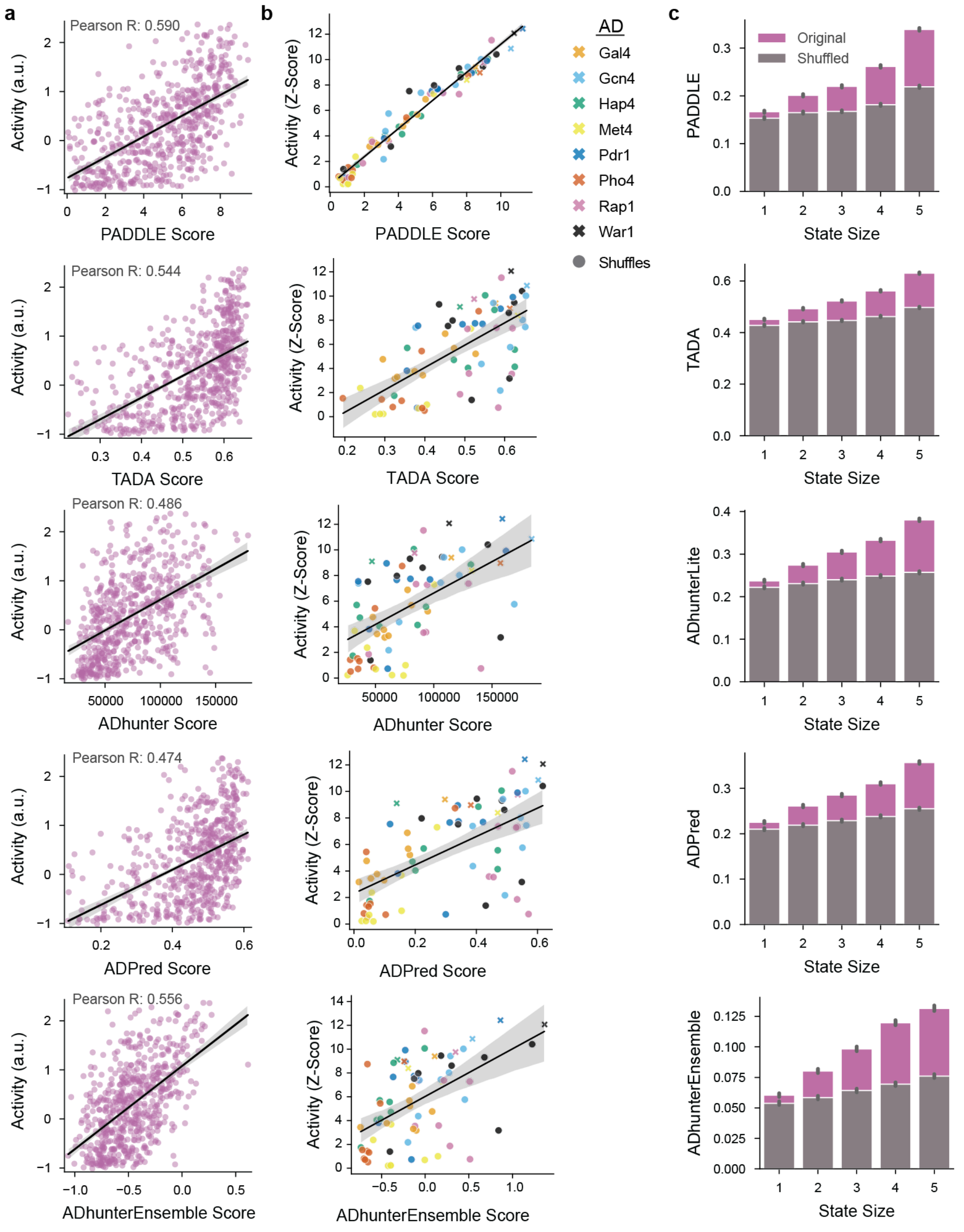


**Extended Data Figure 2. The neural network AD predictors recognize AD grammar.**
a) All neural network models can generally predict the effect of shuffling the Gcn4 AD sequence on activity in the Staller et al. 2018 shuffle mutants. b) All neural network models can generally predict the effect of shuffling sequences on AD strength in the Sanborn et al. 2021 shuffle mutants. Best fit lines are OLS fits with 95% confidence intervals (1000 bootstraps) on the fit. c) Cumulative fraction of 500,000 generated (pink) or shuffled (grey) designed sequences that are predicted by standard thresholds for each AD predictor by state size. Each sequence was shuffled once. Error bars denote 95% confidence intervals determined from 1000 bootstrapping resamples.

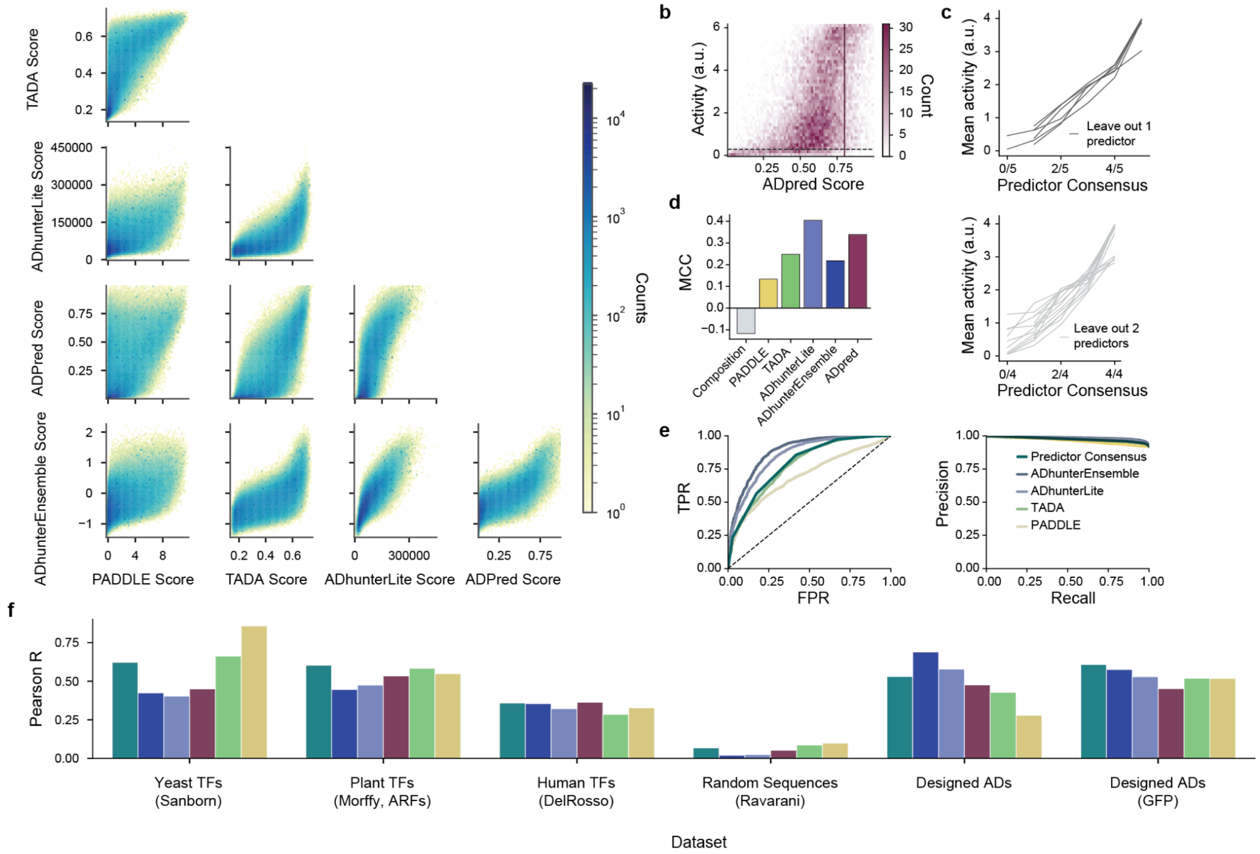


**Extended Data Figure 3. AD predictor analyses.**

a) The neural network AD predictors generate correlated predictions. For five of the neural network AD predictors, we plotted pairwise comparisons of the predictions. b) ADpred average of all 40 per-residue scores per designed AD plotted against measured activity, with threshold denoted by vertical line. c) Mean activity by increasing predictor consensus for all six permutations of leaving out 1 predictor (maximum consensus = 5, top) and all 15 permutations of leaving out 2 predictors (maximum consensus =4, bottom). Trends mirror Figure 4h. d) Matthews correlation coefficient (MCC) calculated for each AD predictor on the tested designed ADs. e) A simple sum of the 6 model predictions (Predictor Consensus) serves as a reasonable classifier of active designed ADs in receiver operator characteristic curves and precision recall curves. f) Pearson R correlation coefficients by dataset for AD predictors and MADRIGAL. Instead of using defined ADs, we used the tiles experimentally tested in the original experiments. See Methods for details on the datasets used.

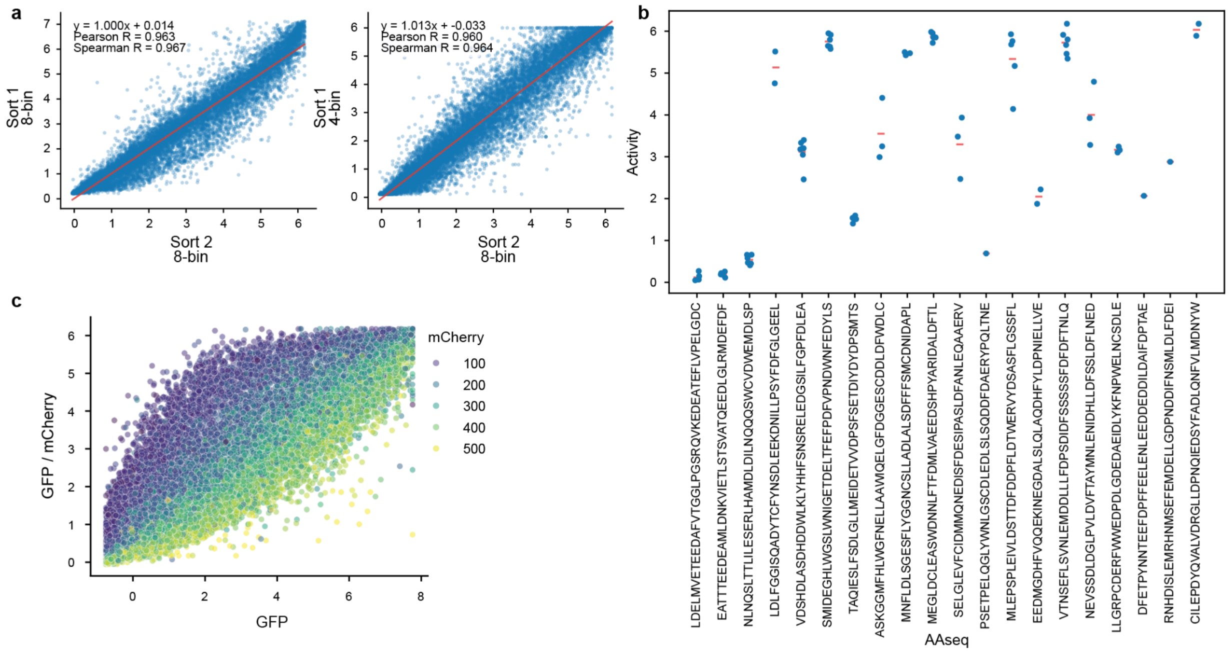


**Extended Data Figure 4: Assessment of Sort-seq measurement quality.**

a) The reproducibility between two independent sorting experiments was higher than our other recently published datasets[1,2]. GFP / mCherry ratio sorting into 4 bins gave consistent results to ratio sorting into 8 bins, even on different days, but 8 bins better resolves the activity of very strong ADs. b) Individual barcodes gave very consistent measurements for 20 randomly selected designed sequences (Sort 2, ratio). Each barcode arises from an independent yeast transformation and is an independent biological replicate. c) Ratio measure of AD activity is highly correlated with the GFP-only measure. X-axis is AD strength measured by GFP-only sort and Y-axis is AD strength measured by GFP/mCherry sort. Sequences are colored by the mCherry measure of TF abundance. For most sequences, the GFP and ratio metrics are highly correlated. For sequences with low mCherry, i.e. low abundance, the ratio metric is higher than the GFP metric, as expected for shrinking the denominator.

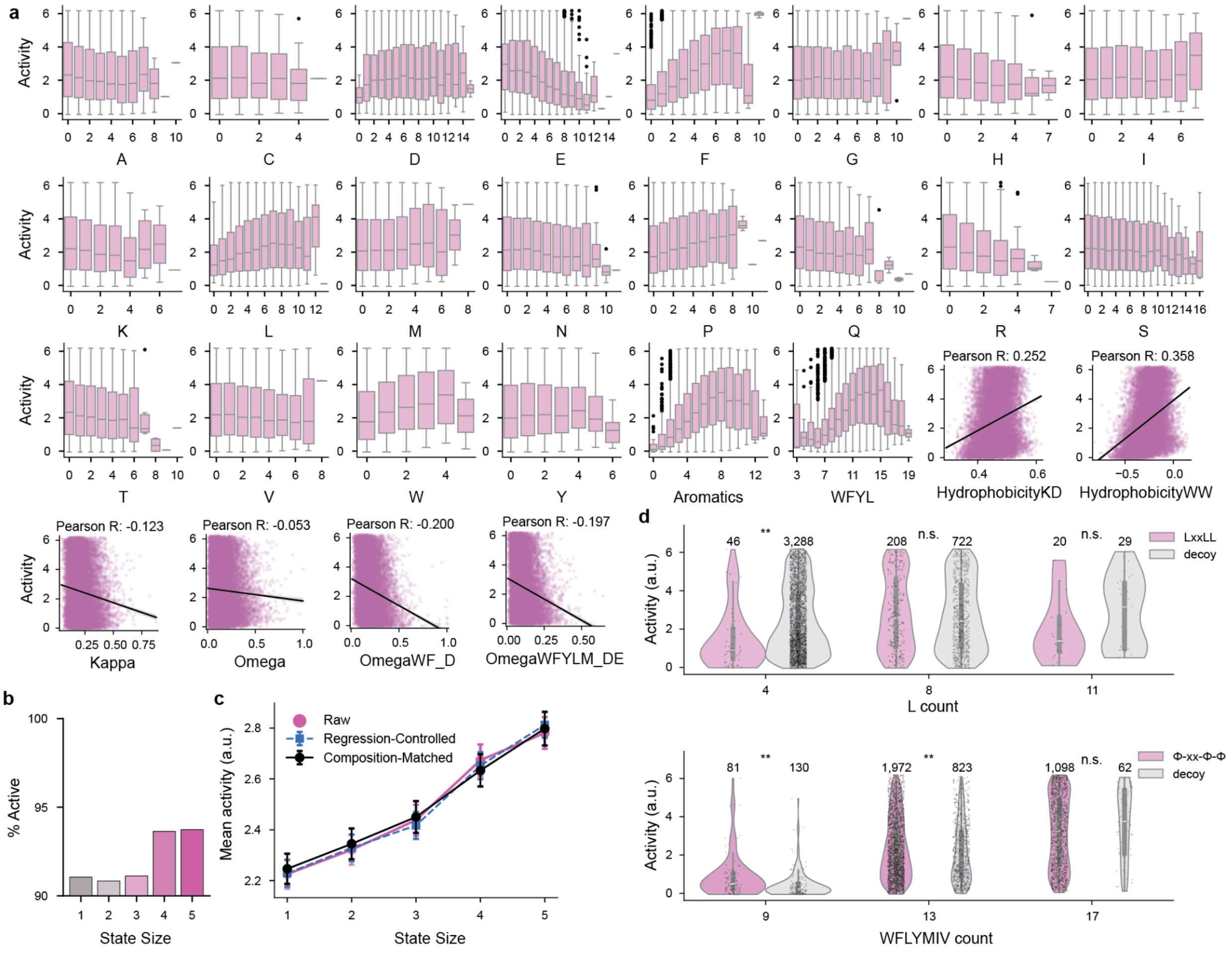


**Extended Data Figure 5. Relationship between composition and AD activity.**

a) For each amino acid, box plots show the relationship between composition and activity. All these trends are consistent with previous observations[2–4]. For the two acidic residues (D,E) the relationship between abundance and AD activity is not straightforward, because the key control variable is the net charge. When combining the count of W,F,Y,L&M residues or of W,F&Y activity rises and falls. The Kyte-Doolittle (KD) and Wimley-White (WW) hydrophobicity scales are shown. The charge patterning grammar signals Kappa and Omega do not show strong relationships with AD activity, but intermixing of acidic residues with key hydrophobic residues was mildly correlated with AD activity[2,5,6]. b) Percent of active designed AD sequences by state size. c) Sequences generated by longer Markov Chains have higher average activity even when we control for composition. "Raw" is the full designed ADs dataset, "Regression-Controlled" is average activity over state sizes after subtracting linear-regression-predicted compositional contributions; "Composition-Matched" measures variation in activity over state size within compositionally similar strata. Error bars are 95% confidence intervals on the mean. d) One motif is associated with increased activity. Distributions of activity for sequences with LxxLL(top) and ΦxxΦΦ (bottom) motifs compared to no-motif sequences with similar motif-relevant residues. Significance evaluated by Welch's two-sided t-test with BH-correction across strata (*q < 0.05, **q<0.01). Φ includes W,F,Y,L,M,I,V.

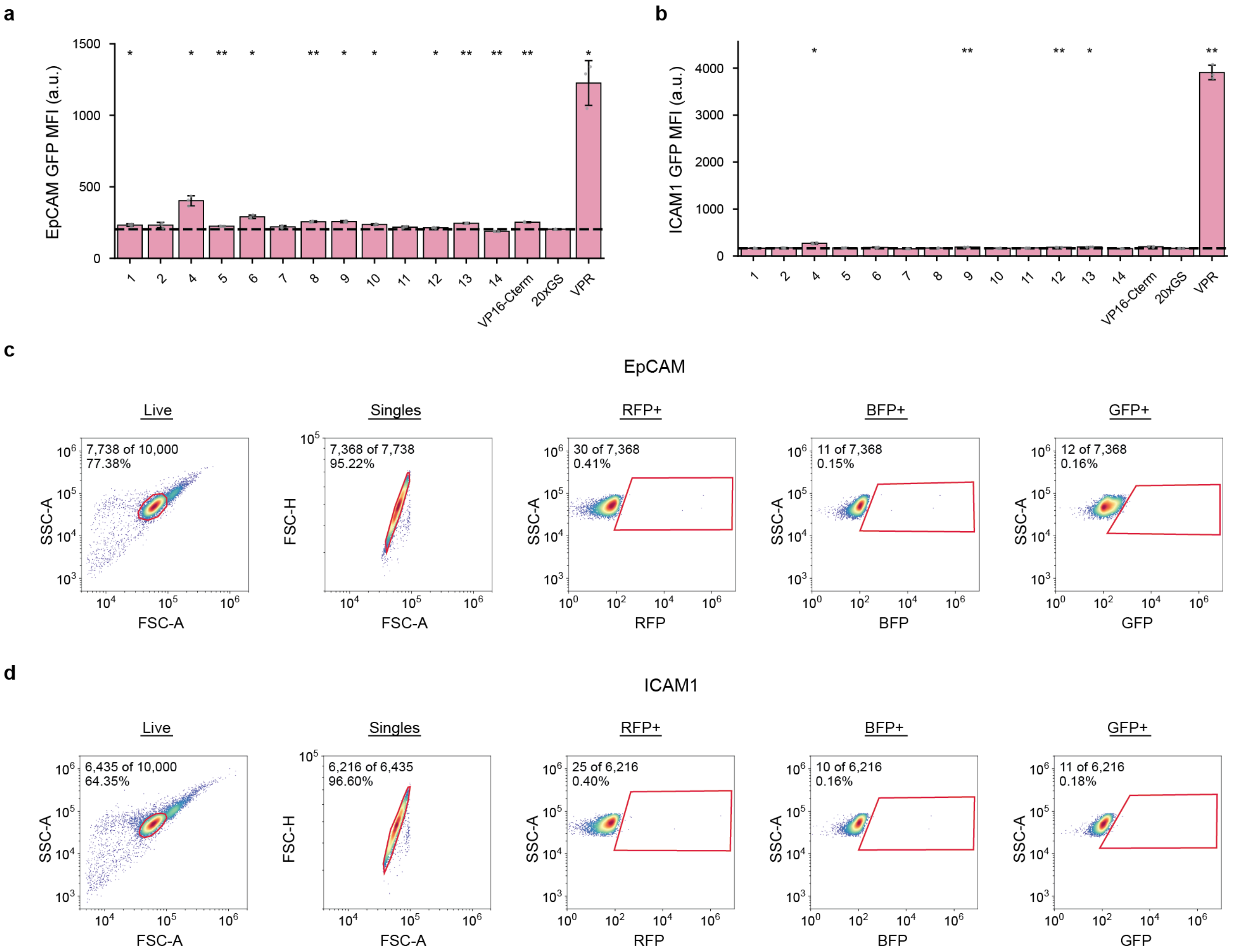


**Extended Data Figure 6. Fluorescence flow cytometry on human cells.**

a) Median fluorescent intensity (MFI) for the GFP signal for the designed ADs tested in HEK293T culture for the EpCAM and (b) ICAM1 surface marker. *q<0.05, **q<0.01, by Welch's two-sided t-test from 20xGS negative control, BH-corrected within each plot. c) Flow cytometry gating scheme on untransfected cells measured with anti-EpCAM and (d) anti-ICAM1 antibodies, respectively. We gated on live cells, then single cells, then transfected cells expressing both (RFP+) and BFP+ cells. GFP+ cells and MFI were quantified from RFP+ and BFP+ cells.

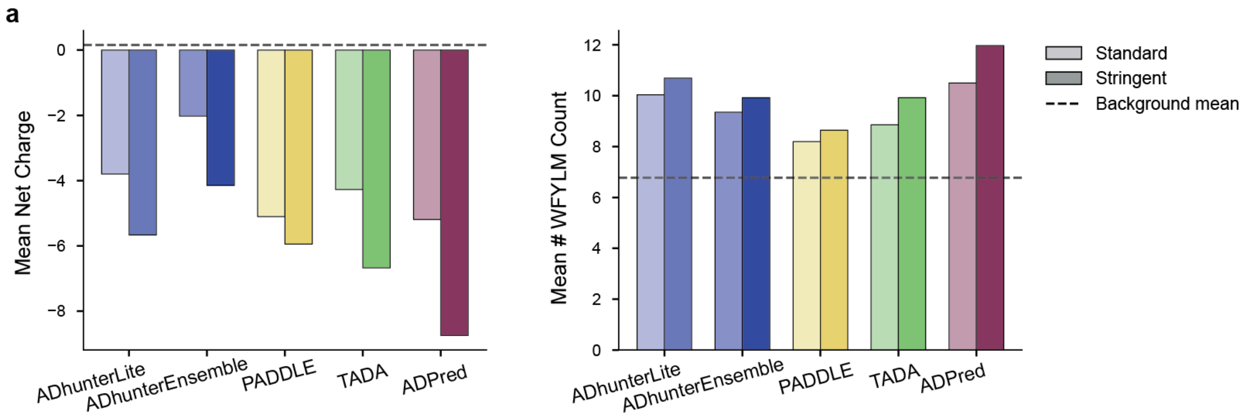


**Extended Data Figure 7. Sequences predicted to be active under more stringent prediction thresholds are more negatively charged and have more hydrophobic residues.**
Using our set of 230,302 sequences, the composition of predicted ADs arising from standard or stringent thresholds was plotted. The dotted line denotes the background mean composition of our sequences.
