## Extended Data Tables for "Machine learning guided design of strong transcriptional activation domains"

List of Supplemental Tables

**Table S1: All designed candidate activation domains**

- Columns: AAseq, ADhunterLite_Score, TADA_Score, PADDLE_Score, ADPred_Score, ADPred, ADhunterEnsemble_Score, Comp, ADhunterLite, TADA, PADDLE (with secondary structure), ADhunterEnsemble, State Size, Comp_original, PADDLE_Score_original (no secondary structure), TADA_Score_original (old version of the model), ADhunterLite_Score_original

**Table S2: Candidate activation domains synthesized for experimental validation**

- Columns: AAseq, FACS_Activity, FACS_StdDev, NumReps_Sort2, Activity_Sort2, StdDev_Sort2, NumReps_GFP, Activity_GFP, StdDev_GFP, NumReps_mCherry, Activity_mCherry, StdDev_mCherry, NumReps_Sort1, Activity_Sort1, StdDev_Sort1, NumReps_Sort1rep, Activity_Sort1rep, StdDev_Sort1rep, State Size, Measured, ADhunterLite_Score, TADA_Score, PADDLE_Score, ADPred_Score, ADPred, ADhunterEnsemble_Score, Comp, ADhunterLite, TADA, PADDLE, ADhunterEnsemble, Comp_original, PADDLE_Score_original, TADA_Score_original, ADhunterLite_Score_original
- The Activity_Sort2 (i.e. the ratio sort from replicate 2) was used for all analyses. It contained more yeast transformants (i.e. more barcodes for each candidate AD) and it contained the Controls Library spiked in.

**Table S3: Controls Library**

- FACS_Activity FACS_StdDev, NumReps_Sort2, Activity_Sort2, StdDev_Sort2, NumReps_GFP, Activity_GFP, StdDev_GFP, NumReps_mCherry, Activity_mCherry, StdDev_mCherry, NumReps_Sort1, Activity_Sort1,StdDev_Sort1, NumReps_Sort1rep, Activity_Sort1rep, StdDev_Sort1rep, NegativeControl
- 200 Negative controls sequences (inactive tiles)
- 100 strong activation domains, positive controls
- 197 activation domains of intermediate strength spanning the range between the negative and positive controls.

**Extended Data Table 1: Linear regression supports an interaction between state size and composition.** Model 1 is linear regression on composition, Model 2 is linear regression on state size, Model 3 is linear regression on both composition + state size, Model 4 is linear regression on composition x state size (their interaction).

| Test | Models compared | Dataset | ΔCV $R^2$ | F | (df) | p |
|---|---|---|---|---|---|---|
| **Composition explains activity** | 1 vs intercept | All tested designed ADs | 0.246 | 290 | 19,16750 | $<10^{-100}$ |
| | | Clustered (50%) tested designed ADs | 0.237 | 248 | 19, 15029 | $<10^{-100}$ |
| | | State sizes 1-4 only | 0.227 | 209 | 19, 13367 | $<10^{-100}$ |
| **State size explains activity** | 2 vs intercept | All tested designed ADs | 0.013 | 226 | 1, 16768 | $<10^{-50}$ |
| | | Clustered (50%) tested designed ADs | 0.008 | 118 | 1, 15047 | $<10^{-26}$ |
| | | State sizes 1-4 only | 0.008 | 114 | 1, 13385 | $<10^{-25}$ |
| **Composition + state size explains activity** | 3 vs intercept | All tested designed ADs | 0.261 | 299 | 20, 16749 | $<10^{-100}$ |
| | | Clustered (50%) tested designed ADs | 0.248 | 250 | 20, 15028 | $<10^{-100}$ |
| | | State sizes 1-4 only | 0.235 | 207 | 20, 13366 | $<10^{-100}$ |
| **Composition x state size explains activity** | 4 vs intercept | All tested designed ADs | 0.267 | 160 | 39, 16730 | $<10^{-100}$ |
| | | Clustered (50%) tested designed ADs | 0.254 | 133 | 39, 15009 | $<10^{-100}$ |
| | | State sizes 1-4 only | 0.237 | 109 | 39, 13347 | $<10^{-100}$ |
| **Adding state size improves on composition alone** | 3 vs 1 | All tested designed ADs | +0.015 | 343 | 1, 16749 | $<10^{-75}$ |
| | | Clustered (50%) tested designed ADs | +0.011 | 223 | 1, 15028 | $<10^{-49}$ |
| | | State sizes 1-4 only | +0.008 | 138 | 1, 13366 | $<10^{-31}$ |
| **Adding interaction terms between state size and composition improves on both** | 4 vs 3 | All tested designed ADs | +0.006 | 10.5 | 19, 16730 | $<10^{-31}$ |
| | | Clustered (50%) tested designed ADs | +0.006 | 8.05 | 19, 15009 | $<10^{-22}$ |
| | | State sizes 1-4 only | +0.002 | 3.94 | 19, 13347 | $<10^{-7}$ |

**Extended Data Table 2: Strong and moderate activation domains selected for follow-up studies in plant leaves and human cell culture.**

| SEQ ID NO | Amino Acid Sequence |
|---|---|
| 1 | KQQEDATSSFPDNDDSSHSLNNSEAVEFEDFLKFSAFLDQ |
| 2 | AELDSDDDFGFCLVPWEMFHLNLVDDWERGQLQTTRSAAS |
| 3 | GELLFDDFGFCYDDKFSSFVNEQEDFEFPELDFLGLEPFS |
| 4 | FDIEDYLDDLDFSSLPPLSESFDLDSLLSGDFFDGLEEFA |
| 5 | DFLDSILKSPDEDPFEPDDFSGFSGGDRFTESLHYYSSLF |
| 6 | EFDDFNFEDYLTFFDETFDINELLGISSSNFLDDCYFSDI |
| 7 | AFTDSPNQLNNILGDEDDPLVPFDQFINLVSDNSNKNAFF |
| 8 | ELTFDEFIEQFTEEDITKLLDTPRSKPSSYFFNYDTSQDY |
| 9 | PPLDPLFLDEDVPLFLDEDVPLFLDEDVPLFLDEFHQSLL |
| 10 | VLEWEEEIDYDWDYHIGEDMLEMMMDTLEGFDLDMIELFG |
| 11 | EEDEFYLDAIASTLGSPTTNSSSSQVGVLPSNLDITFLDF |
| 12 | HYADFFSHDEDAFLLFSPDDNAAGDDLSEFSESENFLFVG |
| 13 | MPLSFKSSDEEDDDDLNMDQDYDFSFLDLTSPTSSLFPPL |
| 14 | ELFDLDFWWFQEQTSKIVLLLVGDGHGLELVPECCSPVEF |

**Extended Data Table 3: Fluorescence values of the bins for the fluorescent activated cell sorting.**

| Bin | Sort2_Ratio | Sort1_Ratio | Sort1_RatioReplicate | Sort1_GFP | Sort1_mCherry |
|---|---|---|---|---|---|
| **1** | 3,212 | 5,815 | 12,176 | 492 | 28 |
| **2** | 12,462 | 17,690 | 40,720 | 1,328 | 165 |
| **3** | 27,384 | 32,080 | 91,331 | 2,316 | 299 |
| **4** | 46,955 | 51,502 | 241,223 | 3,444 | 517 |
| **5** | 75,380 | 81,489 | N/A | 4,916 | N/A |
| **6** | 117,166 | 125,567 | N/A | 6,832 | N/A |
| **7** | 188,575 | 194,220 | N/A | 9,952 | N/A |
| **8** | 262,143 | 262,143 | N/A | 16,581 | N/A |

### Extended Data Table 4: Primers used in this study

| Primer Name | Sequence |
|---|---|
| LC3.P1 | gcgggctctacttcatcggctagc |
| YL_randBCs_R3_3' | AATTCGCTTATTTAGAAGTGGCGCGCCNNNNNNNNNNNNNNCGGGCCCTCAGCTAGTTATCA |
| YP18_cPCR_URA3_US_5’ | CTGCGCCAATTGATGACAATAC |
| CP19.P6 | GACGTTGGTGGAGTCCAAAG |
| YP7_KAN_colPCR_5’ | GATTCTTGTTTTCAAGAACTTGTCATTTG |
| YP19_cPCR_URA3_DS_3’ | ACTTTTCTGAGCACTTCATGATGC |
| CP21.P14 | tcctcatcctctcccacatc |
| CP17.P12 | GGACGAGGCAAGCTAAACAG |
| SL5.F1 | CTTTCCCTACACGACGCTCTTCCGATCTActgcgggctctacttcatc |
| SL5.F2 | CTTTCCCTACACGACGCTCTTCCGATCTTActgcgggctctacttcatc |
| SL5.F3 | CTTTCCCTACACGACGCTCTTCCGATCTCTActgcgggctctacttcatc |
| SL5.F4 | CTTTCCCTACACGACGCTCTTCCGATCTACTActgcgggctctacttcatc |
| SL5.R1 | GTGACTGGAGTTCAGACGTGTGCTCTTCCGATCTTCGCTTATTTAGAAGTGGCG |
| SL5.R2 | GTGACTGGAGTTCAGACGTGTGCTCTTCCGATCTATCGCTTATTTAGAAGTGGCG |
| SL5.R3 | GTGACTGGAGTTCAGACGTGTGCTCTTCCGATCTTATCGCTTATTTAGAAGTGGCG |
| SL5.R4 | GTGACTGGAGTTCAGACGTGTGCTCTTCCGATCTCTATCGCTTATTTAGAAGTGGCG |
| MVS_00XX_I2 | AATGATACGGCGACCACCGAGATCT_index2_ACACTCTTTCCCTACACGACGCT |
| MVS_00XX_I1 | CAAGCAGAAGACGGCATACGAGAT_index1_GTGACTGGAGTTCAGACGTG |
