## Supplemental Note for "Machine learning guided design of strong transcriptional activation domains"

### Supplemental note 1: Comparison of model performance

#### Overview of models

We compare five neural networks and one composition model for predicting ADs from protein sequence. PADDLE, ADhunterLite, and ADhunterEnsemble quantitatively predict AD strength, while TADA and ADpred are classifiers that report confidence scores. There was an additional published neural network we did not include because we could not get it running locally[1]. ADpred and PADDLE are the two first generation neural network models. ADpred is trained on random peptides measured in yeast[2]. PADDLE is trained on fragments of yeast TFs measured in yeast[3]. The Composition Model (Comp) counts the abundance of seven amino acids to predict human activation domains with excellent performance given its extreme simplicity[4]. We have contributed to the development of three second generation neural networks: TADA, ADhunterLite, and ADhunterEnsemble. TADA was trained on sequences from Arabidopsis TFs measured in yeast[5,6]. ADhunterLite (originally called ADhunter_v1[7]) was retrained on fungal Gcn4 homologs measured in yeast[8]. ADhunterEnsemble (originally called ADhunter[7]) was initially trained on sequences from Arabidopsis and yeast measured in yeast, but then improved with active learning on a diverse set of fungal sequences measured in yeast [7]. Predictions between all the models are correlated (**Extended Data Figure 3a**). Despite the diverse origins of these models' training sequences, all these datasets have been measured in baker's yeast with only two different reporter gene promoters, which likely biases the predictions towards acidic and aromatic rich ADs.

An important technical consideration for running the predictors is that they accept different input sequence lengths. ADpred takes inputs of arbitrary lengths and outputs per-residue predictions, using an internal 30-residue sliding window. Here, an AD prediction by ADpred is defined as at least 10 consecutive residues scoring above its threshold of 0.8, as recommended by the authors[2]. PADDLE and ADhunterEnsemble require 53AA, so we append the constant 13AA linker sequence from our plasmid to our designed tiles. We have shown that this heuristic does not cause problems: the first few residues have low importance in sequence analysis[9] and low importance scores when we run model interpreters[7,8]. The original Comp model used 39 AA, but modified the code to accept 40 AA instead. TADA and ADhunterLite natively use 40 AA.

#### Sensitivity to training data and evaluation data

The neural network predictors are extremely sensitive to subtle differences in the AD training task. In this study, we evaluate the Activity of each candidate AD with a Ratio metric: AD activity is the GFP expression normalized to the cellular abundance of the synthetic TF (mCherry expression). We prefer the Ratio metric because it measures specific activity. Erijman et al. and Sanborn et al. use the "GFP output" metric: they define each AD's activity by total GFP expression, without normalizing for TF abundance. Sanborn et al. partially accounted for variation in TF abundance by sorting cells within a narrow range of mCherry signal[3]. We have evidence that this approach does not completely control for variation in TF abundance[8]. The Ratio metric is dominated by the GFP signal, with some modulation by abundance (**Extended Data Figure 4c**). Differences in abundance can influence GFP expression, making the GFP output measurement a convolution of AD abundance with AD strength (**Extended Data Figure 4c**)[8]. AD predictors trained on the Ratio metric (TADA, ADHunter, ADHunterLite) are the top performers by PRC and ROC AUC and Pearson correlation with Ratio data **(Figure 4 b-g**). However, when evaluating on GFP output data, AD predictors trained on the GFP output metric (ADpred and PADDLE) become the top performers (**Figure 5d, Extended Data Figure 3f**). The Comp model, which was designed based on natural human sequences, has greatly improved performance on GFP output data. We conclude that models trained on Ratio perform better on Ratio data while models trained on GFP perform better on GFP data. This subtlety may not surprise some computer scientists but is an important context for users of these models, which are intended for cross-disciplinary use.

**The neural networks can detect local grammar in ADs**

To assess whether the neural networks have learned how sequence grammar controls AD function, we analyzed published data that shuffled the arrangement of residues in several ADs. In Gcn4 shuffling the arrangement of residues leads to sequences that span the full dynamic range of the assay[10]. For 8 53-mer ADs, shuffling generally decreased activity[3]. The Comp model only reads composition so it cannot detect the effect of these shuffle mutations. All five neural network predictors predict the effects of the shuffle mutations well (**Extended Data Figure 2a-b**). This result indicates that the neural networks can detect how sequence grammar modulates AD function.

**Evaluating model performance**

Our goal was to use sequences for which model predictions disagreed to assess model performance. If a sequence was predicted to be active by 2/4 models, this disagreement would mimic uncertainty in an active learning framework [11]. In practice, our final experimentally measured set had few sequences for which the predictors disagreed. This paucity of sequences with disagreement arose for two reasons: 1) we designed the sequences with a high stringency thresholds, but to fairly evaluate the predictors we elected to use the standard, published thresholds for downstream analysis. This effect is particularly apparent for TADA, for which under the published 0.4 threshold, nearly all our experimentally tested sequences are predicted to be active. 2) The model combinations were not evenly sampled because of a coding error in the script that subsampled sequences for ssDNA oligo synthesis (Methods).

Among the existing AD predictors, the worst model is Comp, a simple, interpretable, and mechanistic classifier[4]. Comp likely struggles on designed ADs because it was designed to scan natural TFs, which contain additional sequence context missing in the designed sequences. Comp was so simple we were always surprised it worked at all, so it is reassuring to see that it finally has a limit to its usefulness. The neural network predictors outperform Comp at classifying ADs (**Figure 4g**, **Extended Data Figure 3d**). Sequences in **Figure 4h** predicted by only one model are enriched for being predicted only by the Comp model. This lower success rate for sequences predicted only by the Comp model is because these sequences are called as inactive by all the neural networks, which actually contains a lot of information. Additionally, Comp performance, like PADDLE, improves when evaluating on raw GFP rather than AD activity. Active sequences that are not predicted by Comp tend to be lower abundance, suggesting that the Comp sequence features (W/F/L/D/E/R/K) are also relevant for protein stability [12].

Of the neural networks, PADDLE performed the worst at predicting Ratio data, but it was the best model when predicting raw GFP output (**Figure 5d, Extended Data Figure 3f**). We have some evidence that PADDLE is overfit, because only ⅔ of its top predictions were active in our experiment,[9] and PADDLE performs uncommonly well on its own shuffle datasets relative to Staller et al., 2018 (**Extended Data Figure 2a-b**). Notably, the PADDLE predictions on our designed ADs are more accurate than predictions of systematic mixtures of D's and aromatic residues [13], perhaps because our sequences are more natural-looking. PADDLE is one of the most accurate predictors of human activation domains for unknown reasons[5].

The performance of TADA is difficult to evaluate with the designed sequences. The designed ADs were generated based on the PADI dataset, which was used to train TADA. Likely as a result, TADA predicts almost half of our 500,000 generated sequences as active. There are very few sequences predicted uniquely by TADA (<5% among all generated sequences, <1% of experimental sequences, considering ADhunterLite, PADDLE, and Comp). This absence of negative predictions by TADA makes its performance difficult to assess on the synthetics data. TADA performs well in many scenarios: as a classifier of both ratio activation domain activity and GFP output, and as a regressor, even though this was not its intended function (**Figure 4d**,**g**, **Figure 5d**, **Extended Data Figure 3f**).

ADpred performs very well on the designed sequences. It is a more stringent predictor, because a classifier score of 0.8 is a relatively high-confidence threshold[2]. This might explain why ADpred performs similarly in both ratio activity and GFP output metrics, because the difference between these metrics diminishes as sequences become more active. ADpred was trained on random 30 residue protein sequences, where composition is likely the key driver of activity and natural grammar signals are relatively uncommon.

ADhunterLite and ADhunterEnsemble perform the best on the designed sequences. Both are regressors designed to predict quantitative activity from Ratio data. In addition both were trained using data collected by our group using the same FACS machine, so they may have learned some technical noise associated with this assay. They have also learned specifically to predict ratio activity, and their performance drops when assessed on the GFP output metric (**Figure 5d**, **Extended Data Figure 3f**). The thresholds for both these models are stringent. While thresholding is a useful evaluation metric, ADhunter models are quantitative predictors.

We believe that ADhunterEnsemble is currently the best single model for predicting AD ratio data, but it is computationally intensive to run, requires GPUs, and has inefficient memory usage. We are working to release a more efficient version of ADhunterEnsemble in an update to the original github repository.

#### Analysis of Short linear Motifs

We also searched for Short Linear Motifs (SLiMs) that have been previously implicated in AD activity: WxxLF[14], LxxLL[3,15], and ΦxxΦΦ[16]. WxxLF and LxxLL are present only in 0.4% and 6.6% of tested sequences, respectively, while ΦxxΦΦ is present in 75.4% of sequences. The high hydrophobic content of synthetics means that all three motifs occur at comparable rates by random chance (5.8% and 75.2% for LxxLL and ΦxxΦΦ in shuffled sequences, respectively). The LxxLL motif does not significantly increase activity compared to no-motif sequences with the same hydrophobic content (**Extended Data Figure 5d**). The ΦxxΦΦ boosts activity of sequences with fewer WFYLM residues. This result is consistent with the idea that motifs can boost activity of suboptimal composition[3].
